# *Corylus avellana* non-specific lipid-transfer protein Cor a 8 is a moonlighting enzyme with a new lipase activity

**DOI:** 10.1101/2024.11.22.624807

**Authors:** Fissore Alex, Di Napoli Giulia, De Sciscio Maria Laura, Santoro Valentina, Salladini Edoardo, Marengo Mauro, Oliaro-Bosso Simonetta, Vanzetti Gaia, Caratti Andrea, Dal Piaz Fabrizio, Piazza Francesco, Manzoli Maela, Genova Giuseppe, Barbiroli Alberto Giuseppe, Iametti Stefania, Fraternali Franca, Adinolfi Salvatore

## Abstract

The high-fat content of hazelnuts, mainly triglycerides, makes them prone to lipid oxidation during storage, which has a big impact on their sensory and nutritional quality. The chemical pathways leading to hazelnut oxidative rancidity have been well characterized and it are faster on free fatty acids. Lipase(s) enzymes are required, in oilseed, to hydrolyze the ester bond to free the single molecule of fatty acids. This step, necessary for germination, is the first event to trigger rancidity. Identifying the lipase(s) enzyme and the biochemical pathways involved in rancidity would lead to an effective strategy to prevent fat deterioration. Different proteins have been characterized in hazelnut seed and great interest has been risen towards the non-specific lipid transfer protein family because they were identified as human allergens. Here we show that Cor a 8 – a member of nsLTP – is a novel non-regiospecific lipase that is able to bind to oil-water interfaces and hydrolyze the triacylglycerol (TAGs) ester bonds by a non-canonical active site (non-serine dependent). Molecular modelling and molecular dynamics suggest that Cor a 8 is a moonlighting enzyme not only able to catalyze the hydrolysis of TAGs but also to stabilize the resulting free fatty acids and transport it.

Cor a 8 homologues are present in all land plants, but the specific catalytic amino acids are found only in angiosperms, suggesting an evolutionary adaptation for lipid metabolism unique to flowering plants. This study sets the foundation for understanding this new lipid metabolism in plants and its role in rancidity development.

## Introduction

The European hazelnut (*Corylus avellana* L.) belongs to the *Corylus* genus, Betulaceae birch family, and is one of the 25 existing hazelnut species [1]; originated from the Black Sea region, *C. avellana* L. has been cultivated since Roman times [2].

Hazelnuts are characterized by high-fat content (50-73%), with triacylglycerols (TAGs) as the main constituents of the saponifiable fraction [3], whose major components are the mono- and poly-unsaturated fatty acids (FAs). These FAs are susceptible to autoxidation, leading to the formation of non-volatile primary products (*i.e*., hydroperoxides). These easily decompose to form volatile secondary lipid oxidation products, mainly carbonyl derivatives [4], which lead to rancidity development and impair hazelnut sensory quality.

While the chemistry of rancidity in hazelnuts has been extensively investigated, the biochemical pathways and the enzymes involved have not been fully characterized yet. Indeed, beyond other external factors and autoxidation, lipid rancidity in natural products results from the combination of the enzymatic activity of lipases and lipoxygenases, which could lead to hydrolytic rancidity and oxidation, respectively [4], [5], [6]. Lipases belong to the triacylglycerol ester serine hydrolase family (EC 3.1.1.3), and they do not require any cofactor [7]. Lipases are present in reserve tissues of many oilseed plants and nuts, where they mostly contribute to post-germination oil reserve mobilization, being however not expressed or inactive in the resting and intact seeds [8], [9], [10]. Their activity depends on moisture content and water activity, is independent of oxygen availability, and might not be completely inactivated by roasting [6]. Lipase action results in the release of long-chain FAs, generated by the hydrolysis of tri-, di- and monoacylglycerols [11].

These free fatty acids (FFAs) can be oxidized by lipoxygenases (*enzymatic oxidation*) and/or by autoxidation (*chemical oxidation*) with faster kinetics with respect to those esterified on TAGs. Indeed, recent findings on the evolution of FFAs along with shelf-life in olive oil [12] suggest that post-harvest has a decisive impact on enzymatic activities considering that FFAs are more prone to oxidation than those esterified in TAGs, registering 10 times faster oxidation kinetics [5], [13]. These findings, confirmed also for hazelnuts [4], [6], highlight the relevant role of biochemical pathways able to produce FFAs, in promoting and accelerating chemical oxidation in, but not only, hazelnut seeds [14].

So far, no lipolytic enzymes have been identified in hazelnut seed although several proteins have been well characterized. A large number of oleosins or oil body proteins together with caolesin and steroleosin have been described [15], [16] as well as protease inhibitors and 2S albumin [17]. Moreover, one of the largest and most diversified protein family identified in oilseeds plants are the non-specific lipid transfer proteins (nsLTP) [18].

Plants nsLTPs are usually defined as small (around 9 kDa), basic proteins, widely distributed across all orders of higher plants. Structurally, nsLTPs present four to five α-helices partly wrapped by a long C-terminal segment [19] and contain a conserved motif of eight cysteines, linked by four disulphide bonds, which stabilize a compact α-helix fold forming a tunnel-like cavity for the ligand [20]. This structure confers stability and enhances the ability to bind and transport a variety of hydrophobic molecules. Their highly conserved structural resemblance but low sequence identity reflects the wide variety of ligands they can carry, as well as the broad biological functions to which they are linked to, such as membrane stabilization, cell wall organization and signal transduction [18], [21].

Although many of these proteins have been supposed to have a nitrogen storage function in oilseeds [22], [23], [24], [25], their physiological functions are not fully known yet.

In this frame, the objective of this paper was to identify the hazelnut (*Corylus avellana)* seed lipase(s) and carry out their biochemical, biophysical and enzymatic characterization to elucidate the early stage in seed germination as well as the molecular mechanisms leading to lipid hydrolysis and oxidation when these seeds are stored to be used as key ingredients for the confectionery industry (80% of the production).

Employing a classical biochemical approach, we successfully isolated a new lipase, later identified as Cor a 8 (UniProt Q9ATH2), a well-known nsLTP. Cor a 8 demonstrated hydrolytic activity on both synthetic and natural substrates: 4-nitrophenyl laurate and triolein, respectively. Electron microscopy was used to demonstrate the capacity of the synthetic substrates to form enzyme-bound micelles, a crucial element for distinguishing the newly discovered lipase from an esterase. Notably, Cor a 8 functions as a rare non-specific lipase capable of hydrolyzing the ester bond on both C1 and C2 positions with different affinities.

Subsequently, molecular dynamics techniques were employed to elucidate the interaction between Cor a 8 and its substrates, providing insight into the underlying mechanisms of i) substrate recognition and catalysis on both TAGs C1 and C2 and ii) increased stability of the FFA product while diolein is exiting. These data strongly suggest that the nsLTP Cor a 8 is a new moonlighting non-specific lipase enzyme able to hydrolyze TAGs and also transport single fatty acid chains.

Moreover, it is important to remind that Cor a 8 homologues have been found in all land plants, but the putative catalytic amino acids appear only in angiosperms, suggesting a significant evolutionary adaptation in lipid metabolism of flowering plants.

This study serves as a foundational step towards elucidating the complex processes of lipid metabolism in plants and its ramifications for rancidity development.

## Materials and method

### Protein purification

Commercial samples of raw hazelnuts (*Corylus avellana* L.) were supplied by Soremartec Italia SRL (Alba-CN, Italy), homogenized by a mortar and defatted with hexane 97% by Soxhlet extractor.

For protein extraction, defatted hazelnuts were dried and suspended in a 1:10 ratio (w/v) in 50 mM Tris-HCl, NaCl 100 mM, pH 8.00 (buffer A). After stirring for 2 hours at room temperature, suspensions were filtered and centrifuged (10000 ·g, 30 min, 4°C), and the supernatant underwent anion exchange chromatography on a HiTrap Q-HP column (Cytiva), previously equilibrated with buffer A. After eluting the unbound fraction with buffer A, the bound protein fraction was eluted with 50 mM TRIS HCl, 1 M NaCl, pH 8.00. All fractions were collected and analyzed by Coomassie-stained SDS-PAGE on a 15% gel, followed by enzymatic assay to gauge their lipolytic activity.

Cor a 8 was spectrophotometrically quantified at 205 nm by using a molar extinction coefficient of 282790 M^-1^cm^-1^ [26].

### Microscale lipase assay

Hydrolytic activity of hazelnut extracts and purified proteins on the synthetic substrate 4-nitrophenyl laurate (pNPL) was measured by following on an EnSight® HH3400 multiplate reader (Perkin Elmer) the absorbance increase at 410 nm due to the release of 4-nitrophenol (PNP) at 37 °C, pH 8.00. As a control, an enzyme-free reaction mixture was used.

More specifically, 16 mg of pNPL (Sigma-Aldrich) was dissolved in 2 mL of anhydrous isopropanol. 0.25 mL of the alcoholic substrate stock solution was dissolved in 4.75 mL of Tris 50 mM, NaCl 100 mM, 0.5% (v/v) Triton X-100, pH 8.00, pre-heated to 50°C to reach a pNPL final concentration of 1.24 mM. Self-assembled lipid micelles were used in the enzymatic assays. 0.2 mL-reaction mixtures included 26 µM purified enzyme, preheated at 37°C for 5 minutes, and 0.9 mM substrate, and kinetic release at 37°C of 4-nitrophenol was spectroscopically recorded every 3 minutes.

### Mass spectrometry analysis

High resolution mass spectrometry analysis of the intact purified protein was carried out on an Orbitrap Q-Exactive Classic Mass Spectrometer (Thermo Fisher Scientific) by directly injecting a 100 ng/mL solution of the protein in 0.1% aqueous formic acid:acetonitrile 80:20 (v/v). To achieve protein identification, samples underwent SDS-PAGE and the protein-containing band underwent a trypsin in gel digestion procedure. The peptide mixtures were analyzed on an Orbitrap Q-Exactive Classic Mass Spectrometer (Thermo Fisher Scientific) equipped with a nanospray ion source and coupled with a nanoUltimate3000 UHPLC system (Thermo Fisher Scientific). Peptide separation was performed on a capillary EASY-Spray PepMap column (0.075 mmÅ∼ 50 mm, 2 μm, Thermo Fisher Scientific) using aqueous 0.1% formic acid (A) and CH_3_CN containing 0.1% formic acid (B) as mobile phases and a linear gradient from 3 to 40% of B in 45 minutes and a 300 nL·min^−1^ flow rate. Mass spectra were acquired over an m/z range from 375 to 1500. MS and MS/MS data underwent Mascot software (v2.5, Matrix Science) analysis using the non-redundant Data Bank UniprotKB/Swiss-Prot (Release 2021_03), limited to green plants proteins (Viridiplantae). Parameter sets were: trypsin cleavage; carbamidomethylation of cysteine as a fixed modification and methionine oxidation as a variable modification; a maximum of two missed cleavages; false discovery rate (FDR), calculated by searching the decoy database, 0.05.

### Circular Dichroism (CD)

Circular Dichroism spectra and temperature ramps were recorded in a Jasco J-810 spectropolarimeter equipped with a Jasco PFD-425S Peltier system, by using a 0.1 mm path length quartz cuvette. Protein was dissolved 0.1 mg/mL in Tris-HCl 5 mM, pH 8.00, adding when specified 2 mM DTT. The samples were subjected to an analysis cycle including an initial spectrum, a heating temperature ramp, a hot spectrum, a cooling temperature ramp, and a final spectrum. Spectra were recorded from 190 to 260 nm (scanning speed 10 nm/min, data pitch 0.1 nm, response 0.5 sec, band width 1 nm). Temperature ramps were recorded between 20 to 90°C, monitoring the CD signal at 222 nm (temperature slope 1°C/min, data pitch 0.1°C, response 2 sec, band width 1 nm). CD signal was normalized in term of mean residue ellipticity (MRE). the α -helical content was estimated from the MRE at 222 nm [27].

### FESEM combined with EDS analysis

Field Emission Scanning Electron Microscopy (FESEM) analyses were performed by a Tescan S9000G FESEM 3010 microscope (Tescan Orsay Holding a. s., Brno- Kohoutovice, Czech Republic) working at 30 kV, equipped with a high brightness Schottky emitter and fitted with Energy Dispersive X-ray Spectroscopy (EDS) analysis by an Ultim Max Silicon Drift Detector (SDD, Oxford, UK). For the analysis, one drop of the prepared samples was deposited onto an aluminum stub coated with a conducting adhesive and subsequently left to dry in the air at room temperature. The dried samples were submitted to Cr metallization (ca. 5 nm) using an Emitech K575X sputter coater (Quorumtech, Laughton, East Sussex, UK) to avoid charging effects and then inserted into the chamber by a motorized procedure. Images were acquired by using the secondary electrons (SE) detector by in-beam high resolution mode and in-beam cross-free mode. Histograms of the particle size distribution were obtained by considering a representative number of particles observed in the images for each examined sample: The mean particle diameter (d_m_) was calculated through the following equation:

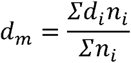

where n_i_ the number of particles with diameter d_i_.

By this approach, not size-selected lipid micelles composed by pNPL and Triton X-100 were analyzed in the presence of 26 µM enzyme. Control measurements were performed on: I) not size-selected lipid micelles composed of Triton X-100 (0.5 %) and pNPL (0.4 mg/mL); II) not size selected lipid micelles composed of pNPL 0.4 mg/mL and enzyme 26µM; III) a Triton X-100 (0.5%) solution with added enzyme (26 µM); IV) a solution of enzyme (26 µM) alone.

### Thin Layer Chromatography

25 µg of triolein dissolved in dichloromethane were added to 100 µL of isopropanol containing Triton X-100 5% and evaporated under nitrogen. The dried micelles were dissolved in 100 μL of buffer A and 70 µM of purified lipase was added and diluted to a final volume of 500 μL. The mixture was incubated overnight at room temperature with gentle shaking. The hydrolytic reaction products were extracted twice with 1.5 mL of diethyl ether. Then, 700 µL of extract was evaporated, dissolved in a small amount of dichloromethane CH2Cl2 and spotted on TLC plates (Alufolien Kieselgel 60F254; Merck, Darmstadt, Germany) using n-hexane/ diethyl ether/acetic acid ethyl acetate (70:30:1; v/v) as developing solvent. The spots were made visible in iodine vapor or by UV. Triolein, 1,2-diolein, 1,3-diolein, and oleic acid were used as standards.

### Analysis and fits of the enzyme kinetics

Experimental kinetic curves for the Cor a 8 enzyme were obtained by measuring the formation of product over time at different substrate concentrations. A stock substrate solution was prepared as previously described. Each substrate concentration analyzed was obtained diluting the stock solution with Buffer A and 0.5% triton-X to keep constant the percentage of triton in each assay. 1:1 lipase-substrate mixture was used to conduct the assay.

In the absence of any specific microscopic information, the simplest, thermodynamically consistent kinetic scheme for enzyme catalysis is the reversible Michaelis-Menten (RMM) scheme. In order to fit the kinetic model (see supplementary material) to our data, we developed a fitting strategy to minimize a global cost function, defined as the total least-squares deviation between the measured and predicted product concentrations computed over all time points and all considered substrate concentrations. For this purpose, we developed a routine integrates numerically the RMM rate equations in correspondence of a given point in parameter space and used a minimization strategy based on the Differential Evolution (DE) method [28] to find the global minimum of the cost function (S5). DE is an evolutionary fitting algorithm suitable for solving complex global optimization problems via an iterative evolutionary process. Since it is intrinsically stochastic in nature and does not require initial guesses for the free parameters (only bounds), the result of many independent optimization runs can be used to assess the statistical variability of the best-fit results. In Figure supplementary 5 it can be seen that all the best-fit values are distributed around their means (those reported in Table 2) with a spread, measured in units of standard deviation over mean, that is at most 4 %.

### Computational analysis and molecular docking

Docking simulations with pNPL were conducted using the HADDOCK 2.4 webserver, based on the shape-restrained docking protocol [29]. We used the crystallographic structure of maize nsLTP (PDB: 1FK0) [30] as a reference, and docked the small molecule NPC12 to the hazelnut Cor a 8 structure (PDB: 4XUW) [31]. A total of 64 NPC12 conformers were generated [32], and 200 of the best rigid-body models were refined.

The final models were clustered with an RMSD cut-off of 1.5 Å.

Since the Cor a 8 crystal structure (PDB: 4XUW) is in an apo form with a closed lid, we remodelled the loop residues (98-110) using Modeller [33] to obtain an open form, which was then used for docking with triolein (TOG). The TOG structure was retrieved from literature, and docking was performed using AutoDock Vina [34], [35]. Binding poses were evaluated based on Vina scores and the proximity of polar amino acids to TOG’s polar heads [36]. The three best poses were further refined with short 200 ns Molecular Dynamics (MD) simulations.

For diolein (DOGL) and oleic acid (OLE) systems, the structure was derived from the most sampled conformer during the TOG MD simulations and transforming the 1,2-diolein and the oleic acid into their hydrolysis products. All-atom MD simulations of the complexes were carried out with Gromacs [37] and CHARMM36 force field [38]. The systems were centred in a periodic box, solvated with TIP3P water, and neutralized with ions. Multiple equilibration steps were performed, gradually increasing temperature [39] and decreasing position restraints, before production runs of 1 µs for protein-TOG and 1.5 µs for protein-DOGL-OLE [35]. The simulations used the LINCS algorithm for bond constraints and PME for electrostatics [40], [41], with analysis performed using Gromacs and MDTraj [42].

### Phylogenetic analysis

The Cor a 8 protein sequence served as bait for conducting searches in sequence databases via BLAST. The selected UniProt IDs with the corresponding species are described in Table S1. Multiple alignments were performed with the ClustalW method with all settings set as default [43] and visualized using the WebLogo tool [29]. The tree was constructed by using the program Jalview [30] removing the signal peptides and using a BLOSUM method. Finally, the tree was visualized in the program iTOL [31] where the branch lengths were arbitrary.

## Results

### Lipolytic enzyme(s) purification

The setup of a successful purification protocol for an enzyme requires a suitable and reliable method to measure its activity. Lipase hydrolytic activity was assessed on the synthetic substrate pNPL, which allows a direct spectrophotometric detection of the lipolytic activity product, namely 4-nitrophenol. This method enabled the identification of the fractions containing the enzyme of interest at each purification step. All kinetics measurements were recorded at 410 nm, and the released 4-nitrophenol was quantified by using an extinction coefficient of 18.3 mM^-1^cm^-1^.

Crude proteins extraction from defatted hazelnuts was performed under continuous gentle rotation for two hours at room temperature, by adding TRIS-HCl 50 mM, NaCl 100 mM, pH 8.00 (buffer A) to the defatted crushed hazelnut. This procedure ensured complete extraction with a good protein yield. To isolate the enzyme(s) with lipolytic activity, the extract was loaded onto an anion exchange chromatographic column (HiTrap Q-HP column, Cytiva). An early protein peak (unbound) was observed during low saline elution (Figure 1A, peak 1) while a second protein peak, bound to the stationary phase (bound), was eluted by increasing the ionic strength (Figure 1A, peak 2). Both fractions (unbound and bound) as well as the crude protein extract were analyzed for their hydrolytic activity onto pNPL to identify the fraction enriched with lipolytic enzyme(s).

**Figure 1.**
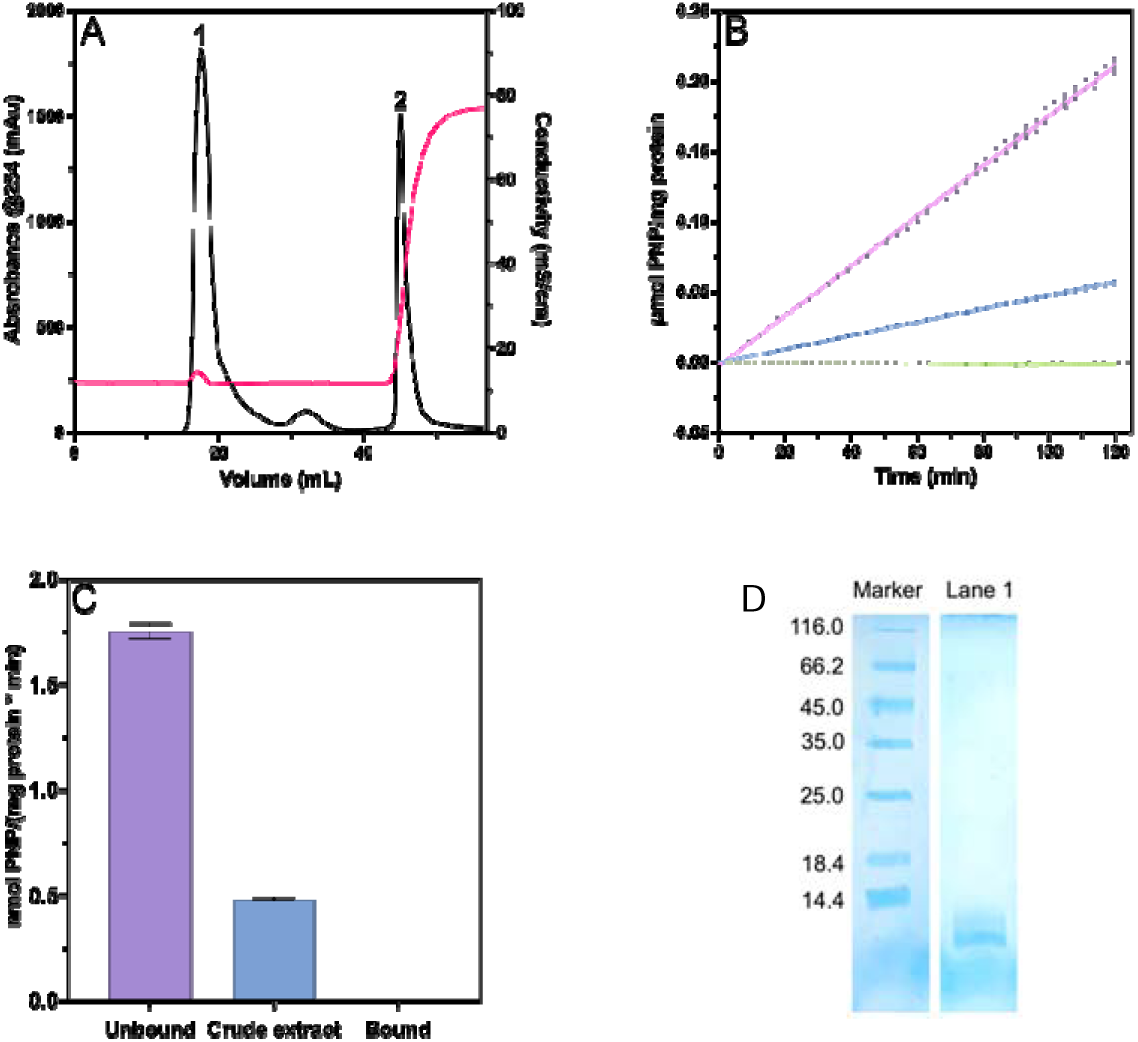
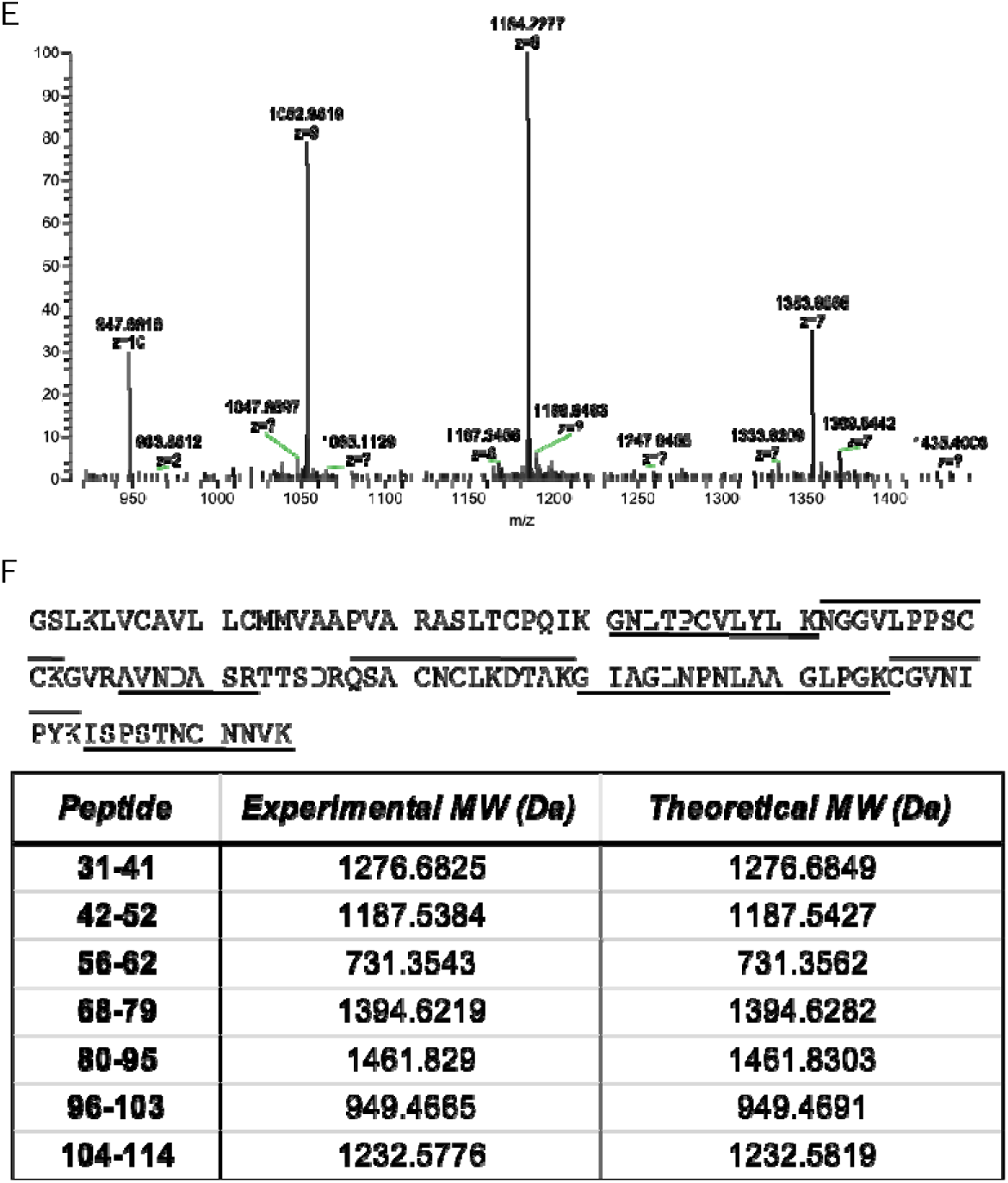
Lipase purification and mass spectrometry-based identification. A) Anion exchange chromatography using a Cytiva HiTrap™ Q HP column. A large fraction of the proteins (unbound fraction) was eluted in a major peak (peak 1). The bound fraction (peak 2) was eluted after application of 1 M NaCl isocratic flow, (red line is the conductivity measured during the chromatography). B) Lipase activity of different fractions of protein during the anion exchange purification. Kinetic measurement was followed using 4-nitrophenyl laurate on crude hazelnut extract (light blue), unbound fraction (purple) and bound fraction (pink). C) Initial rate of crude extract and unbound fraction have been calculated from the kinetics in Figure 1B. The unbound peak (Purple bar) shows a 4-fold increase in lipase activity with respect to the crude lysate (Light blue bar). D) SDS-PAGE to assess the purification protocol: Line M, Proteins as molecular weight marker, Lane 1, unbound peak eluted by the anionic exchange chromatography. E) High resolution ESI-MS spectrum of the intact protein showed a molecular weight of 9465.7 Da. F) The LC-MS/MS analysis of the tryptic digest of the protein allowed the identification of Cor a 8 but the observed peptides covered only 66% of the protein sequence. The difference in molecular weight (theoretical 11806.1 Da) of Car a 8 and the incomplete sequence coverage can be attributed to the cleavage of an N-terminus signal peptide from the mature protein. The cleaved N-terminus is not detected in the ESI-MS of the intact protein, nor is represented in the tryptic peptides. Indeed, the first tryptic peptide detected starts at aa 31.

A moderate formation of 4-nitrophenol was observed in the total hazelnuts extract (Figure 1B, light blue line) while the unbound fraction showed significant lipolytic activity (Figure 1B, purple line). Conversely, the bound fraction showed a negligible activity (Figure 1B, green line). As illustrated in Figure 1C, the unbound fraction (purple bar) exhibited a considerable 4-fold increase of enzymatic activity with respect to the crude extract (light blue bar). These results indicate that most of the lipase enzyme(s) were eluted within the unbound fraction.

The efficiency of lipase purification was confirmed by SDS-PAGE, where a single homogenous Coomassie-stained protein band (Figure 1D, lane 1) was identified in the enzymatically active unbound fraction obtained from the anion exchange chromatography, indicating that the purification protocol was successful.

The apparent molecular weight of this single protein in the unbound fraction was determined via size exclusion chromatography and found to be around 9250 Da (Figure S1), consistent with a monomeric state of the enzyme. It is noteworthy that all lipases characterized so far show molecular weights between 19 and 60 kDa) [44]. This surprising result suggests the possibility of a new lipase with a low molecular weight.

### Identification of the new hazelnut lipolytic enzyme: Cor a 8

It was necessary to identify the new purified enzyme. Mass spectrometry (LC-MS) was used, and a single protein was found in the purified sample with an experimental MW of 9465.7 Da (Figure 1E). The protein was identified by subjecting it to in-gel trypsin digestion followed by high-resolution mass spectrometry analysis of the resulting peptides. The bioinformatic elaboration of the data led to the identification of Cor a 8, with a 66% coverage of the full sequence (Figure 1F). The missing fragments were all located at the N-terminus of the protein, where the presence of a secretion peptide composed by the first 23 amino acids was previously described. The lack of the N-terminal region in the purified mature protein was also consistent with the difference observed between the experimental and the theoretical (11806.1 Da) molecular weight of Cor a 8. Remarkably, also the results of the peptide mapping analysis confirmed the presence of this only protein, thus confirming the efficacy of the purification procedure.

Importantly, Cor a 8, one of the most studied allergen in *C. avellana* and classified as a non-specific lipid transfer protein (nsLTP), has been here identified unambiguously as a lipolytic enzyme

### Secondary structure characterization and thermostability

The newly identified Cor a 8 lipase secondary structures and stability were analyzed by circular dichroism (CD). Cor a 8 spectrum shows the typical shape of a structure dominated by alpha helices (two minima at 208 and 222 nm; Figure 2A, blue line), as already reported in literature [32], with an alpha-helical content corresponding to 58.4%. In temperature ramp experiments the protein underwent progressive denaturation, where the unfolding ensemble was not completely reached at 90°C (Figure 2B black line). Indeed, the lipase retained an alpha-helical content of 31.6% at 90°C (Figure 2A red line).

**Figure 2.**
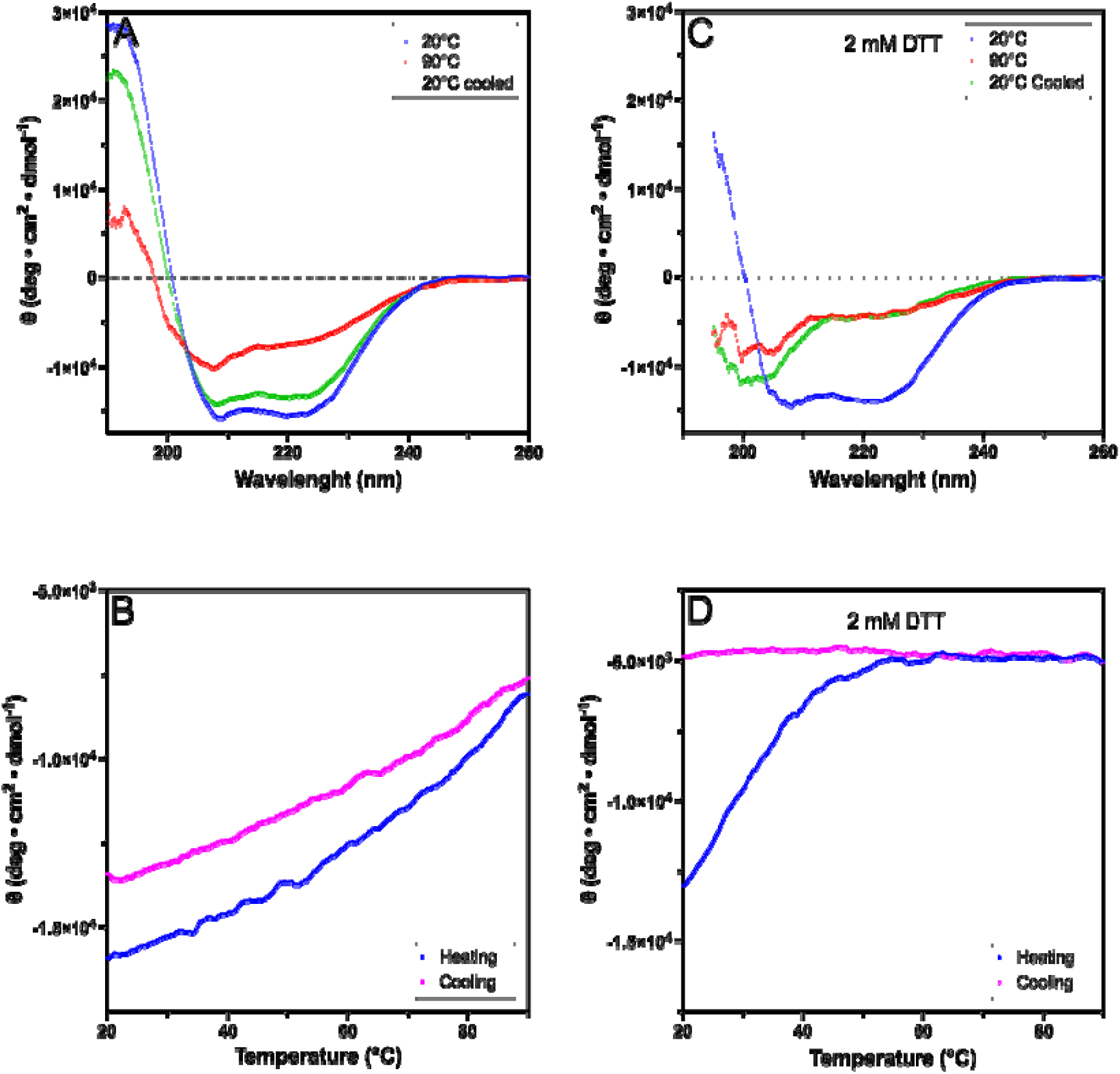
Secondary structural analysis of Cor a 8 by circular dichroism (CD) spectroscopy. A) far-ultraviolet (UV) CD spectra of Cor a 8 at 20°C (blue), after thermal unfolding at 90°C (red) and after cooling it back down at 20°C (green). B) thermal denaturation curves of Cor a 8 monitored by far-UV CD. The melting curve (blue) and the refolding curve (magenta) were obtained by recording the signal at 220 nm as a function of temperature. C) far-ultraviolet (UV) CD spectra of Cor a 8 when DTT is added at 20°C (blue), after thermal unfolding at 90 C (red) and after cooling it back down at 20°C (green). D) thermal denaturation curves of Cor a 8, when DTT, is added monitored by far-UV CD. The melting curve (blue) and the refolding curve (magenta) were obtained by recording the signal at 220 nm as a function of temperature

The unfolding was mostly reversible, as demonstrated by the refolding trace shown in Figure 2B (magenta line) and by re-recording the spectrum after cooling at 20° C (Figure 2A green line). Indeed, the refolded lipase presented 51.4% alpha-helical content, which differs only slightly from the value of the native enzyme. This indicates solubility and lack of aggregation at high temperature. These results agree with other nsLTPs [33].

The high thermal stability could be due to the presence of the four disulfide bridges. To verify this hypothesis, the lipase thermal unfolding was carried out in the presence of 2 mM reducing agent (DTT). In this condition, the lipase still presented a well-preserved signal for the secondary structures (Figure 2C, blue line) comparable to the one without DTT (Figure 2A blue line). On the other hand, thermal stability was highly compromised, reaching the complete unfolding around 60°C as shown by the curve in Figure 2D (blue line). The lipase denaturation in the presence of DTT was irreversible (Figure 2D magenta line), as confirmed by CD spectrum after cooling (Figure 2C green line). Indeed, the lipase almost loses the alpha-helical content at 90°C in presence of DTT (Figure 2C red line).

### Cor a 8 is present at the oil-water interface: Strong indication of being a lipase

Esterases and lipases have previously been distinguished on the basis of their substrate specificity [34]. Lipases require an oil-water interface for their activity, while esterases cannot catalyze hydrolysis in these conditions [45]. Demonstrating that the pNPL used as a substrate can form micelles that mimic an oil-water interface would considerably strengthen the claim that Cor a 8 is a novel active lipase in the hazelnut kernel.

FESEM combined with EDS mapping allowed us to visualize, for the first time, lipid micelles formed by pNPL and stabilized by the presence of Triton-X-100. These micelles provide the oil-water interface required for a lipase enzyme activity. This approach allowed us to gauge the elemental composition of the samples close to the surface of the lipid micelles and to assess both morphology and location at the nanometric level of the enzyme on the lipid micelles.

Despite the poor contrast, due to the low beam current and voltage employed to avoid modification or damage of the sample during the observation, size-heterogeneous lipid micelles of pNPL stabilized by Triton X-100 with rounded shape and d_m_= 670 ± 395 nm were observed (Figure S2). Conversely, a completely different morphology, i.e. a residual organic matrix embedding the buffer salts particles coming from the evaporation of the sample solution, was noticed in the case of the sample containing the enzyme with the lipid micelles formed by pNPL substrate alone. This means that the presence of Triton X-100 is required to stabilize the lipid micelles (Figure S3). Figure 3A and C show that the lipid micelles have spherical shape and size between 250 and 500 nm, with an average diameter of 408 ± 226 nm (Figure 3B). EDS analysis indicates that the main constituent elements of the sample are C, O, N and S, consistent with the composition of lipase (Figure 3D-F). In this case, the lipid micelles were clearly observed, indicating that the presence of the enzyme makes the micelles more stable under the electron beam. Indeed, the enzyme itself is stable (Figure S4). Interestingly, the lipid micelles, in the presence of the enzyme, displayed a coating not observed without it (respectively, Figure 3A and Figure S2), demonstrating that they were indeed covered by the enzyme. This was confirmed by the EDS mapping, which returns a higher relative N amount (14.30 atomic %) with respect to the bare lipid micelles stabilized by Triton X-100 and in the presence of pNPL (spectrum 1, 3.19 atomic %), as shown in Figure 3E, 3F and in Figure S2.

**Figure 3.**
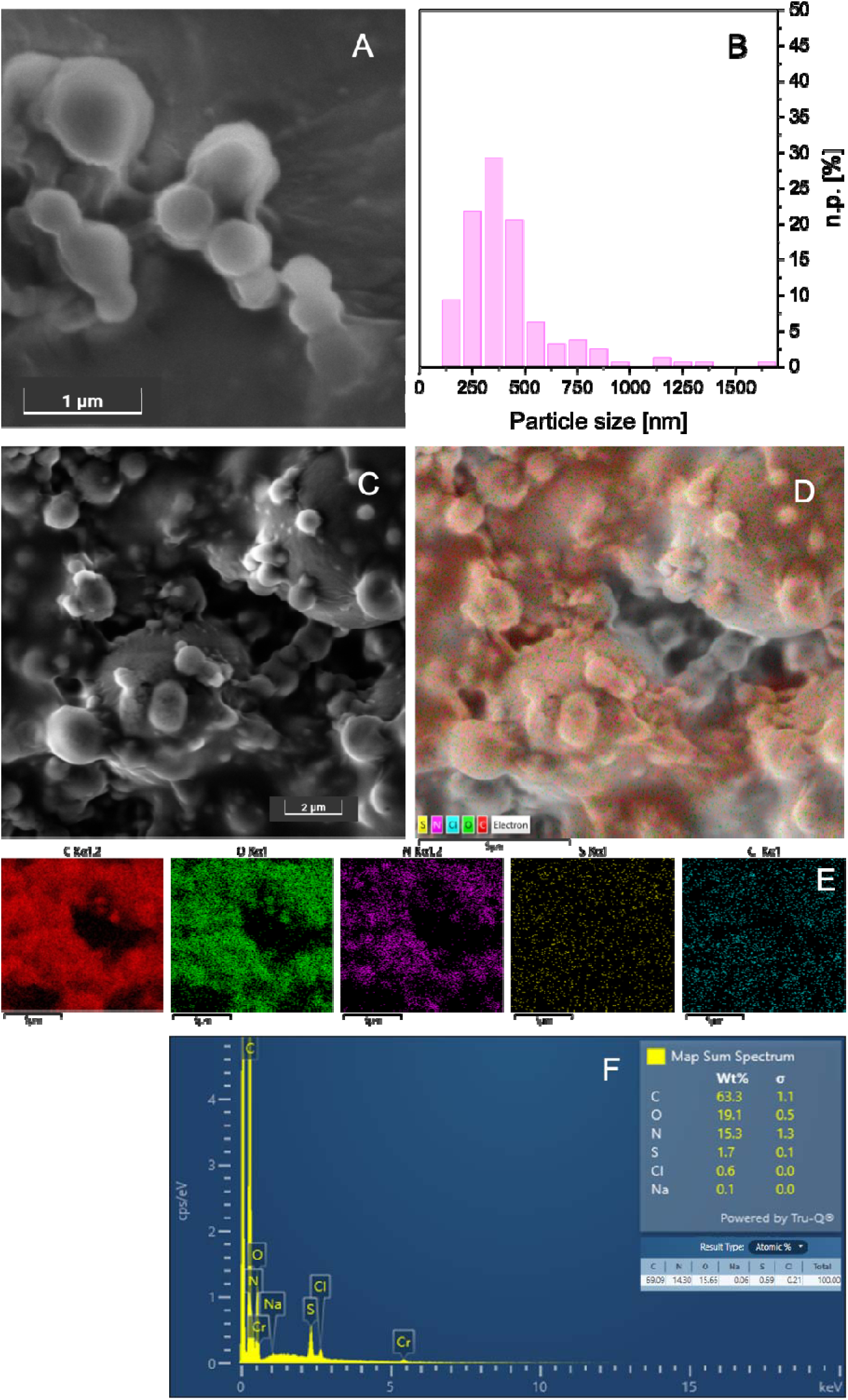
Field Emission Scanning Electron Microscopy (FESEM) of Cor a 8 and pNL micelles. . FESEM representative images (A and C) and particle size distribution (B) of the Cor a 8 on the lipid micelles stabilized by Triton X-100 and in the presence of the pNPL substrate. EDS layered image (D), EDS maps of the region shown in b) for C, O, N, S and Cl (E) and EDS map sum spectrum (F). Images collected at 5 keV with the standard SE detector. n.p.[%] represents the number of counted particles of diameter di. Instrumental magnification: 50000× and 20000×, respectively.

Therefore, according to the amount of N measured through the EDS analysis, it was proven that the micelle outer surface is enriched in N, indicating that the lipase molecules sit preferentially at the surface of the lipid micelles.

These results strongly suggest that the enzyme behaves as a proper lipase, demonstrating its tell-tale ability to physically sit at the oil-water interface.

### Cor a 8 is a new lipase

To further strengthen the claim that Cor a 8 is a new lipase identified in the hazelnut seed, the hydrolytic activity of the purified enzyme was tested on triolein (TOG), one of the most relevant natural substrate, which is the most abundant TAG in the hazelnut seed. It is important to highlight that TOG is not soluble in aqueous solution but tend to form micelles [35], [36]. An overnight incubation of Cor a 8 with triolein micelles stabilized by Triton-X was carried out and the lipidic part of this reaction was subsequently analyzed by TLC (Figure 4, lane 4). Triolein micelles stabilized by 1% Triton-X without the enzyme (Figure 4A lane 2) and micelles composed of triton-X without triolein with purified enzyme were incubated in the same conditions (Figure 4A lane 3). A standard mixture containing triolein, oleic acid, 1,2-diolein and 1,3- diolein was used. In Figure 4A, only in Lane 4 it is possible to observe a significant decrease of TOG, whereas new spots appeared, corresponding to oleic acid, 1,2 and 1,3-diolein, with a prevalence of 1,2- diolein. The presence of both 1,2 and 1,3 diolein strongly suggests that Cor a 8 is a rare, non-specific lipolytic enzyme [37].

**Figure 4.**
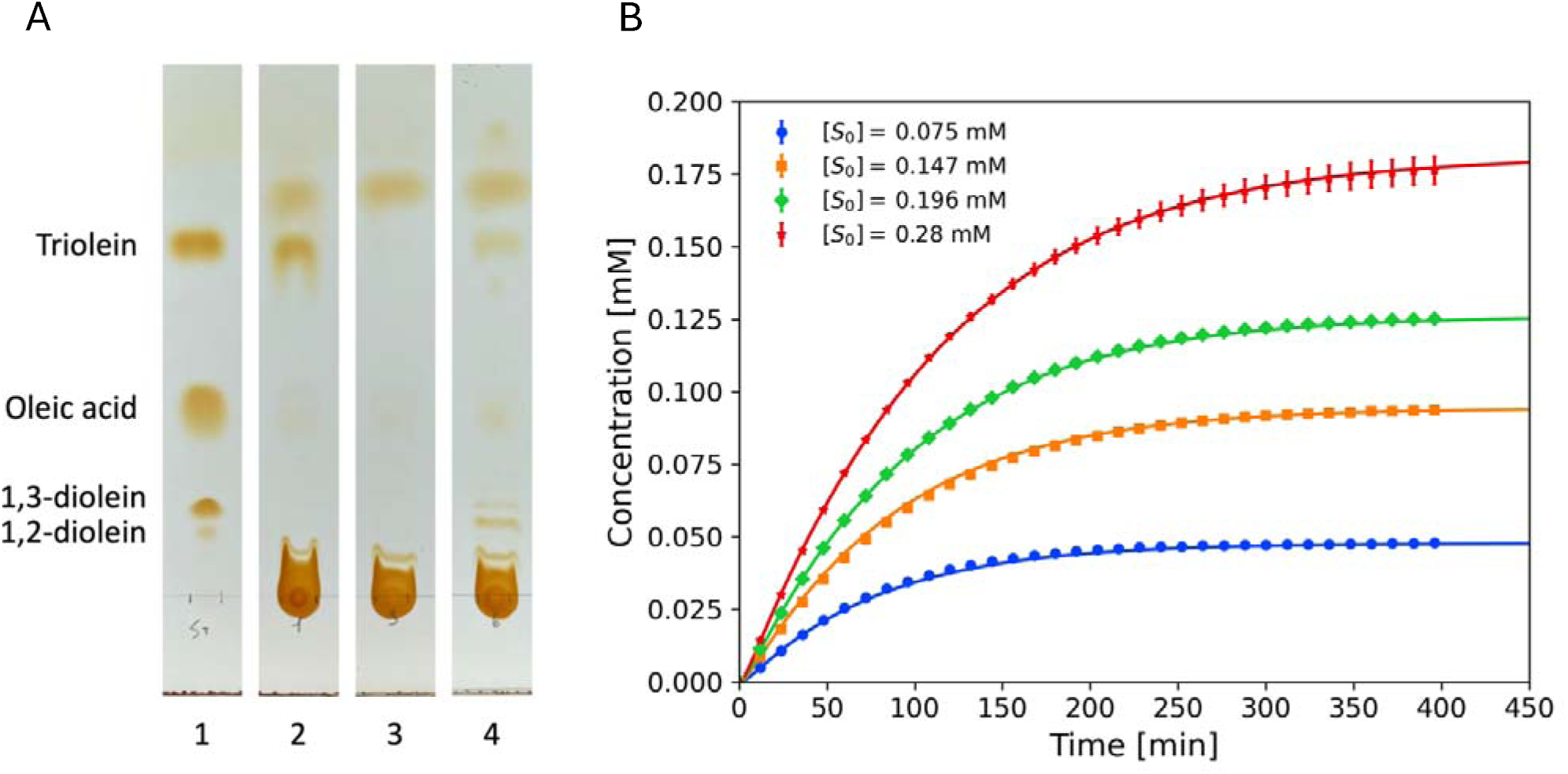
Cor a 8 Lipase activity on triolein and its kinetics measurement. A) thin layer chromatography. Lane 1 – Standards of triolein, oleic acid, 1,2-diolein, and 1,3. Lane 2 – 3 triolein substrate and purified enzyme as control of the enzymatic reaction respectively. Lane 4 - Cor a 8 overnight reaction with triolein micelles. Three new bands of the triolein hydrolysis products are visible corresponding to 1,2-diolein, 1,3-diolein and oleic acid. B) Measured product concentration for the kinetics of the purified enzyme at four different values of the initial substrate concentration [*S*_0_] (symbols) and best-fit solution obtained by numerical integration of the reversible MM equations (solid lines). The average values of the MM best-fit parameters, as well as the corresponding MM constants, are reported in Table 2. Table 1, average best-fit values of the free parameters (first six lines) computed over *N_R_* = 400 independent minimization runs of the stochastic minimization algorithm (see text and supplementary material). The statistical errors on the mean are calculated from the computed standard deviations as 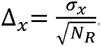. The last two lines report the corresponding average values and statistical errors of the Michaelis-Menten parameters obtained from the fit.

**Table 1.**
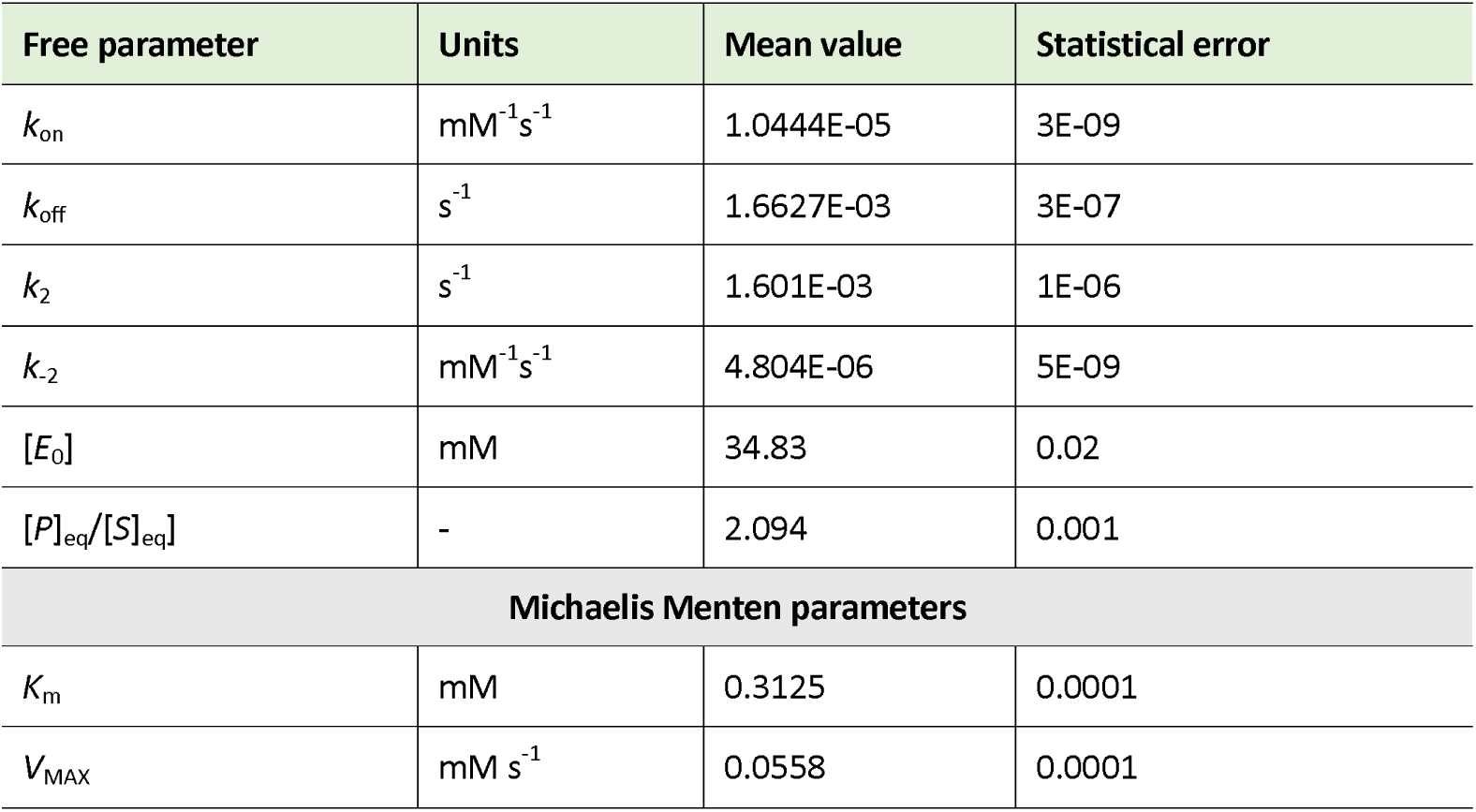

Lipases can be classified as regiospecific or non-regiospecific based on the types of diacylglycerols produced during catalysis. Cor a 8 is identified as non-regiospecific, as it generates both 1,2 and 1,3-sn-diacylglycerols, showing a preference for the 1 acyl position over the sn 2 position, indeed characteristic of non-regiospecific lipases.

### Kinetics characterization

The purified enzyme kinetics was studied under the simplest assumption for the underlying kinetic mechanism, namely a reversible Michaelis Menten (RMM) scheme. Typically, initial slopes are calculated from the measured product concentration at short times and plotted against the initial substrate concentration [S0] (MM plot) to determine the maximum velocity (Vmax) and apparent affinity (Km).

However, restricting to the initial slopes does not guarantee that the catalysis is described by any particular kinetic scheme, as in most cases the product concentration is linear in time at short times. Instead, one should seek to fit the full-time evolution of the product to a given kinetic model with a single set of kinetic parameters for all values of [S0]. Therefore, we simultaneously fitted the reversible MM differential equations to time-resolved kinetic data recorded at all substrate concentrations (see Methods and Figure S5).

The results reported in Figure 4B demonstrate excellent agreement with the prediction of a reversible MM model, with a predicted affinity Km of 0.312 mM, consistent with literature values (table 1) [34].

Thanks to the simultaneous fit of the time course of product generation at all substrate concentrations, our approach tightly constrains the theory to all experimental data in a single, robust fitting algorithm. In particular, this allows for the active enzyme concentration [E]_0_ to be treated as a fitting parameter. Its best-fit prediction turned out to be around 34.8 mM (table 1), a value just 2 times lower than the face value concentration, confirming the soundness of our theoretical approach. Additionally, the equilibrium product and substrate concentrations exhibit a ratio of approximately 2.1 (table 1). Since the predicted value of [E]_0_ is much lower than both [P]_eq_ and [S]_eq_, it can be safely assumed that their sum is quasi-conserved, that is, [P]_eq_ + [S]_eq_ ≈ [S]_O_. This allows us to estimate the fraction of substrate consumed as *f_S_* = 1 - [S]_eq_ ⁄[S]_O_ = [P]_eq_⁄([P]_eq_ + [S]_eq_). Therefore, our measured value [P]_eq_ /[S]_eq_ ≈ 2.1 (see table 2) translates immediately to J_S_ ≈ 0.68. Hence, about 68 % of the initially present substrate molecules appear to be hydrolized by the enzyme It is important to stress that the theoretical model did not consider explicitly the nature of the enzyme-substrate interaction. In fact, in the MM scheme the substrate is treated as freely diffusing, whereas we know that it is only present at the surface of the nano-micelles (Figure 3 panel A). Hence, our best-fit value of the association rate *k*_on_ should be treated as the 3D equivalent of the true 2D association rate.

Incorporating the true binding geometry into the model is a complex task, due to uncertainties in characterizing the surface coverage of substrate molecules on the micelles. Hence, comparisons between different lipases should only be made between data obtained under strictly identical conditions.

### Cor a 8 is a moonlighting enzyme

The discovery of Cor a 8’s hydrolytic activity raises the possibility that this protein might function both as a catalyst and a non-specific lipid transporter. To investigate these moonlighting properties, we conducted a molecular-docking study of Cor a 8 with the synthetic substrate pNPL. This simulation provided a detailed structure of the Cor a 8-pNPL complex, offering insights into its enzymatic reaction. Given the high sequence identity between Cor a 8 and maize nsLTP [32], the crystallographic structure of the latter was used as a reference [38]. The simulation revealed that the hydrophobic tails of the ligands are nested within the hydrophobic cavity, while the head groups (4-nitrophenol) extend outward, interacting with surface residues and solvent molecules (Figure S6A). Figure S6B shows the residues involved in these hydrophobic interactions, suggesting that proper occupation of the hydrophobic cavity is key for substrate binding. The same simulation was carried out using the crystallographic structure of the Peach nsLTP (*P. persica*) and produced similar results.

To further explore Cor a 8’s newly identified lipase activity, we performed Molecular Dynamics (MD) simulations using triolein (TOG), its natural substrate. Despite the absence of a typical catalytic triad (Ser, His, Asp) in Cor a 8, we identified residues potentially involved in binding and catalysis. We also examined the protein’s selectivity towards specific carbons (C1, C2, and C3) in the hydrolysis mechanism. Given the experimental indistinguishability of positions 1 and 3 in solution, our focus was on positions C1 and C2, which correspond to the formation of 1,2-diolein and 1,3-diolein, respectively. Using a modeled open form of Cor a 8, docking experiments generated a Cor a 8-TOG complex where the TOG’s polar head is primarily coordinated with Tyr103. Five 1 µs MD simulations showed that the lipid remains inside the protein, demonstrating Cor a 8’s ability to accommodate a lipid molecule the size of TOG, aided by the flexibility of the binding loop (Figure 5). The hydrolysis process likely requires at least one water molecule near the site, with polar side chains in the active site region either restraining the ligand or assisting in forming a bonded enzyme-ligand complex.

**Figure 5.**
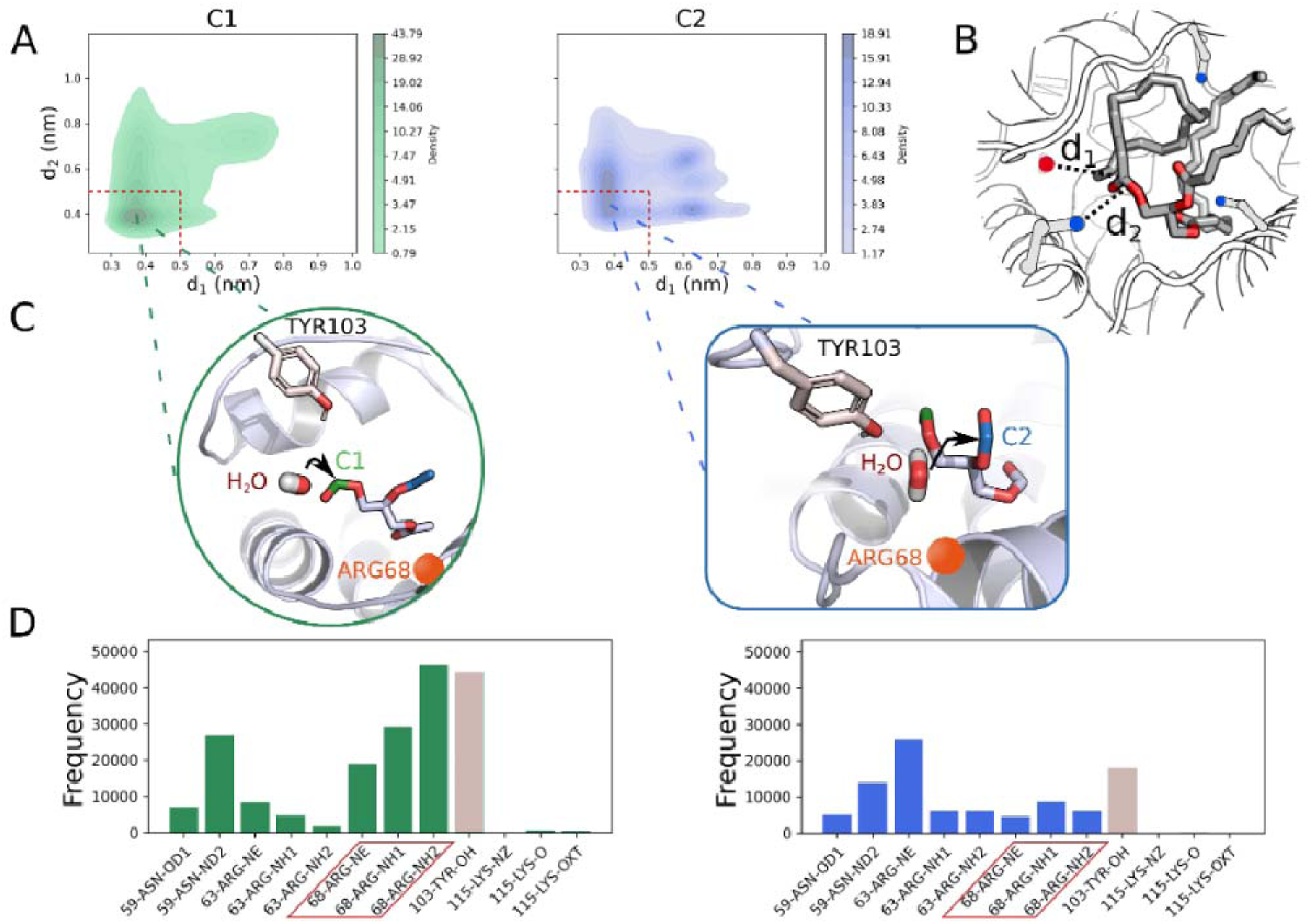
Molecular Dynamics simulations shedding light on the triolein (TOG) selective hydrolysis mechanism. A) Kernel Density Estimation (KDE) of TOG reactive space, as defined by the minimum distance between each carbon atom (available to potential water attack) of the lipid and the oxygen atom of neighbouring water molecules (d_1_, x axis) and the minimum distance of the same carbon atom and the polar atoms of neighbouring residues’ sidechains (d_2_, y axis). The reactive area, i.e. where both distances are lower than of 0.5 nm, is delimited by the red dashed lines. B) Pictorial representation of the descriptors (d_1_ and d_2_, dashed black lines) used to define the reactive space. The protein is shown in cartoon while the lipid is in sticks. Solid blue points represent polar atoms (i.e. both N and O) on sidechains while drawn balls (red for oxygen and white for hydrogens) figure a water molecule. C) Main ‘reactive-like’ conformers, extracted by cluster analysis on all the structures satisfying the reactive-like criterion (under the dashed line in KDE plots), for C1 (left, green) and C2 (right, blue). TOG polar head is shown as sticks in grey while the atoms on the first and second oleic chains are in green and blue. Tyr103, one of the most interacting residues, is shown as sticks while Arg68 position is highlighted with an orange sphere. C) Frequency of observed interactions for each subset of reactive-like conformations between the residues composing a putative “active-site” (selected as the residues within a sphere of 0.6 nm centred on C1 and C2 of the most sampled conformation) and the two TOGL carbon atoms that can be hydrolysed. Zoom out of atoms shown in panel C, Arg68 and Tyr103, are highlighted with a red box in the first case, while the bar color is grey for the latter residue.

To understand the binding modes of selected lipids and their hydrolysis selectivity, we defined a "reactive space" around each TOG ester bond (C1 and C2), focusing on regions where a carbon atom (C1 or C2) has both a nearby water molecule and a polar side chain atom. We measured distances between these C atoms and water oxygens (d1), as well as between the C atoms and polar side chain atoms (d2) and used Kernel Density Estimation (KDE) to analyze these interactions (Figure 5A and B).

Our analysis revealed a single reactive minimum for C1, located entirely within the defined reactive space, suggesting C1 is the preferred site for hydrolysis. In contrast, three minima were identified for C2, but the main minimum was shallower and only partially within the reactive zone, indicating a lower likelihood of hydrolysis at C2 (Figure 5A). These findings align with experimental data, which show a preference for hydrolysis at C1, leading to the formation of 1,2-diolein over 1,3-diolein.

Figure 5C illustrates the binding modes of C1 and C2 reactive complexes, showing that both are similar in structure. Key residues, particularly Tyr103 and Arg68 (Figure 5C and D), play a critical role in the hydrolysis process by forming hydrogen bonds with the ester group and nearby water molecules. These interactions help lock the group to be hydrolysed in a reactive conformation. Additionally, Arg63 and Asn59 contribute to a hydrogen bond network that stabilizes the interactions between key residues, water molecules, and the lipid’s polar head. Notably, C1 interactions are more stable and long-lasting compared to C2, as evidenced by the frequency of contacts between C1 and the active site residues.

To further investigate the structural dynamics of Cor a 8 during hydrolysis, we conducted additional MD simulations with the hydrolysis products, 1,2-diolein and oleic acid. Principal Component Analysis (PCA) was used to simplify the analysis by focusing on the protein’s Cα atoms and the heavy atoms of the lipids. By projecting the sampled conformations onto the essential subspace defined by the first two eigenvectors (which accounted for approximately 70% of the total motion variance), we generated a free energy surface [39] (Figure 6A).

**Figure 6.**
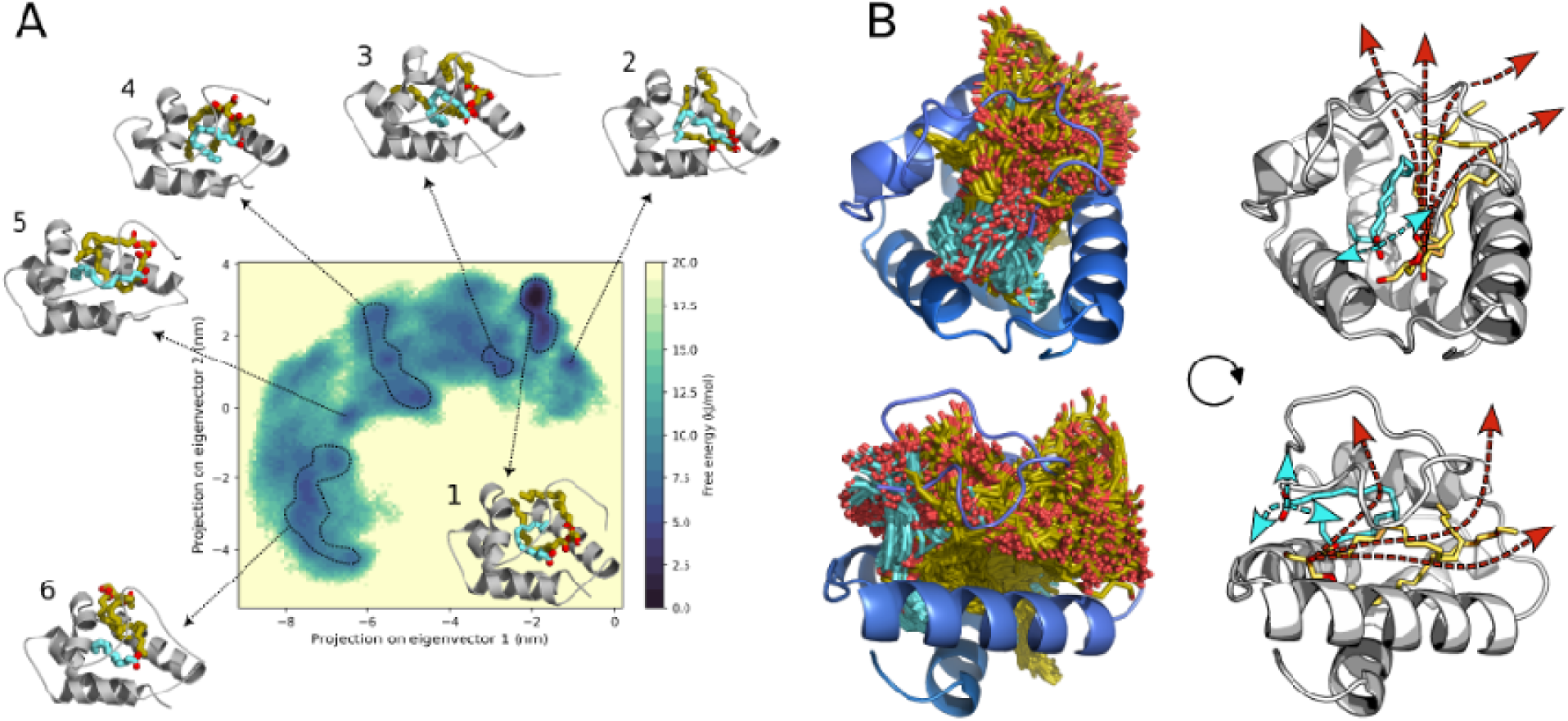
Preferred (un)binding sites for the hydrolysis products, oleic acid and 1,2-diolein, as obtained by MD simulations. A) Free energy landscape of protein-products complex defined by the first two Principal Components (representing around 70% of the total variance). A representative structure of the complex is reported for each subset of conformations. The two ligands are shown in sticks; oleic acid is cyan while diolein is yellow. B) Superposition of randomly extracted frames of the complex in front view (upper panel) and after a rotation of 45° of both y and z axis (bottom panel). For clarity, just the initial conformation of the protein is shown in cartoon while all the sampled conformations of the ligands are in sticks (oleic acid in cyan and the diolein in yellow). On the left, the arrows (cyan for OLE and yellow for DOGL) show the main motions of the ligands’ polar heads with respect to the starting structure, shown as sticks (ligands) and cartoon

This analysis showed that the most stable conformation, represented by the lowest free energy minimum (snapshot 1 in Figure 6A), closely resembles the starting structure, where both lipids remain within the protein’s core. Other conformations in the PC1-PC2 plot reflect the displacement of 1,2-diolein from the initial conformation, with several pathways leading to its unbinding from the enzyme’s cavity. Conversely, oleic acid consistently moves towards the core center, forming stable interactions with Tyr103 and Arg68. These residues form a "gate" that prevents oleic acid from exiting the cavity, ensuring its stable coordination within the protein core.

The different stability of the two lipids is further highlighted in Figure 6B, where a superposition of randomly selected conformations from the MD simulations shows a greater displacement of 1,2-diolein’s polar heads compared to those of oleic acid. Although complete unbinding of 1,2-diolein is rare in the simulated conditions and time frame, the frequent partial unbinding events observed in all simulations suggest that Cor a 8 functions as a moonlighting enzyme. It not only catalyses the hydrolysis reaction but also selectively transports the single-chain lipid, releasing the complementary part.

### The new lipase is conserved in all land plants

NsLTPs are ubiquitously present throughout the plant kingdom. According to Fleury et al. 2019, they show high evolutionary divergence, but the structural fold is conserved, suggesting that they could share a common, yet unknown function [40].

To explore the evolutionary trajectory of nsLTPs, sequences ranging from non-vascular land plants (such as liverworts and mosses) to angiosperms were aligned, revealing variations in sequence identity among different plant groups. These variations hint at possible functional divergence among non-vascular land plants, ferns, gymnosperms, and angiosperms over evolutionary time. Specifically, Figures 7A and S7 highlight the conserved amino acids through sequence logos.

**Figure 7.**
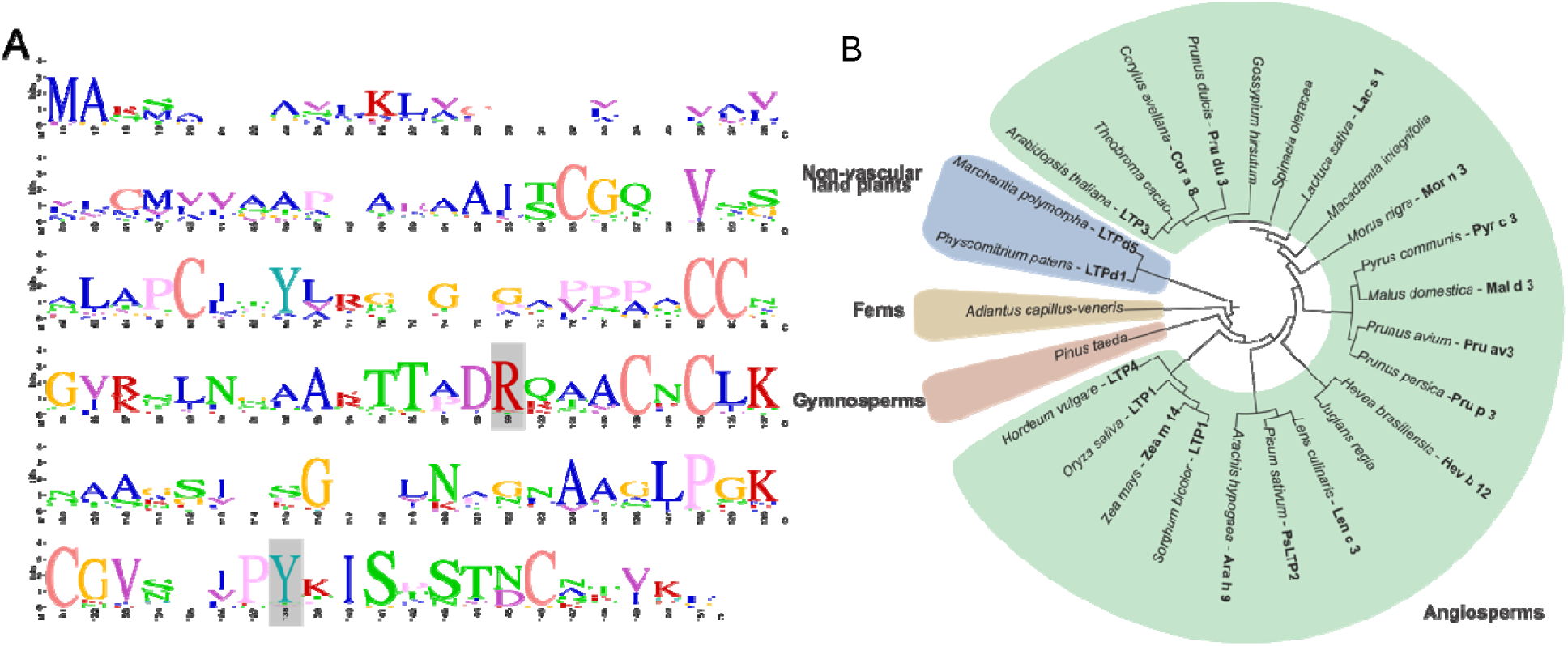
Multiple sequence of Cor a 8 homologues and phylogenetic tree. A) Sequence logos illustrate the conservation of the nsLTP protein, excluding the signal peptide. The residues are marked according to the clustal colour scheme and the Cora8 corresponding Arg68 and Tyr103 are shown as grey sticks. B) Circular phylogenetic tree of nsLTP protein sequences displayed with the scientific name and, where present, the protein/allergen name. Branch lengths are arbitrary.

A critical observation from the alignments is the conservation of the eight-Cys motif, essential for stabilizing secondary structures via four disulfide bridges. This motif is consistently preserved across different plant species, underlining its importance in maintaining the structural integrity of nsLTPs (Figure 7A).

Moreover, two highly interacting residues identified in this study—Arg68 and Tyr103—are notably conserved across many species, although some discrepancies are observed in ferns, mosses, pinus, and liverworts (Figure 7A). These residues have been highlighted by molecular dynamics simulations as key reactive centers, forming hydrogen bonds with the substrate’s ester group and a nearby water molecule. This interaction suggests that Arg68 and Tyr103 play a crucial role in the enzyme’s catalytic mechanism. The phylogenetic tree derived from these alignments (shown in Figure 7B) provides further evidence to support the hypothesis that nsLTPs originated early in plant evolution. Furthermore, the absence of critical enzymatic residues in early land plants suggests that the newly identified catalytic function may have developed more recently in angiosperm. This finding implies a functional evolution in nsLTPs, with the catalytic role emerging later than the initial appearance of these proteins in early land plants.

## Discussion

The chemistry of rancidity, a major concern for the food industry’s need of long-term raw material storage, have been extensively investigated [41], [46], whereas the associated biochemical pathways have been poorly studied in most oilseed plants like *C. avellana*.

Rancidity arises from the oxidation of long-chain fatty acids [12], and is significantly accelerated when these are free, unesterified from triacylglycerols (TAGs) [5]. The oxidation process involves lipases, enzymes responsible for hydrolyzing the ester bonds in TAGs. Given the potential impact on food storage, there is growing interest in identifying, purifying, and characterizing the specific lipase responsible for this process in *C. avellana* oilseeds.

To achieve this, we resorted to a classical biochemistry approach, where the hydrolysis of a synthetic substrates 4-nitrophenyl laurate was used to identify lipolytic enzymes in each purification step. We managed to obtain a purified protein around 9 kDa that was later identified as Cor a 8, the well-studied allergen belonging to the non-specific lipid transfer protein (nsLTP) family. This is a diverse group of low-molecular weight proteins present in all land plants with low sequence homology but a well conserved conformational structure and a basic isoelectric point (between 9 and 11).

All nsLTP structures present a hydrophobic cavity, highlighting their ability to bind and transport various hydrophobic molecules, particularly fatty acids [47], [48] This feature is crucial for their involvement in many biological functions such as membrane stabilization, cell wall organization, and signal transduction, as observed in *in vitro* studies [18].

Our surprising results show that Cor a 8 presents a clear catalytic activity, characterized by kinetic parameters that are typical of lipases. This suggests that Cor a 8 is a moonlighting enzyme able to hydrolyze the ester bond between the fatty acid chain and glycerol as well as to stabilize and transport the resulting FFA where needed.

The discovery that Cor a 8 could be more than just a lipid transporter, potentially functioning as a moonlighting enzyme capable of hydrolyzing ester bonds, was surprising. To substantiate this hypothesis, electron microscopy (FESEM) was employed to demonstrate that the synthetic 4-nitrophenyl laurate can form micelles mimicking an oil-water surface. This was a crucial step towards classifying Cor a 8 as a new lipase, along with its ability to hydrolyze triolein (a plausible natural substrate). Previous studies have explored how different 4-nitrophenyl derivatives with long fatty acid chains can form micelles using techniques like light scattering and DLS [49]. However, we were able to gather richer information by means of carefully conducted electron microscopy experiments (FESEM). This powerful approach revealed that 4- nitrophenyl laurate micelles are clearly coated by Cor a 8 when the protein is added. This is the expected behavior for a lipase, which localizes at the surface of lipid micelles. The ability of the 4-nitrophenyl laurate to form micelles and the ability of Cor a 8 to hydrolyze triolein are strong evidences that Cor a 8 is a new lipase.

Interestingly, Cor a 8 does not present either the serine or the canonical catalytic amino acid triad typical of serine hydrolases. Our molecular dynamics simulations carried out in presence of triolein helped us identify two residues that are well-conserved along the phylogenesis of the nsLTP protein family (Tyr103, Arg63), which mainly coordinated TOG’s polar head in the simulations.

The products released by Cor a 8 are oleic acid and either 1,2 diolein or 1,3 diolein. This evidence led us to conclude that Cor a 8 is a non-specific lipase able to hydrolyze the ester bond in both carbon sn1 and 2.

Incidentally, this rare lipase without apparent regio-preferences could be a potential biocatalysts for the production of high-value TAGs via esterification/transesterification and biodiesel via alcoholysis of TAGs [50], [51].

In order to clarify how lipids bind to the enzyme and the selectivity in the hydrolysis process, a "reactive space" was defined, based on the MD simulations sampled conformations, for each potential TOG ester bond (C1 and C2). The structural features of the C1 and C2 reactive complexes showed similarities, with residues Tyr103 and Arg68 interacting the most with the ester group, forming crucial hydrogen bonds with it and a nearby water molecule to facilitate the hydrolysis process. However, the C1 reactive conformations are more likely to occur and, once attained, the interactions with the neighboring residues are more persistent compared to the C2 ones.

Further MD simulations revealed that within the enzyme’s hydrophobic core, the 1,2-diolein tends to unbind more readily than the oleic acid, which moves deeper into the core and interacts with Tyr103 and Arg68. These interactions form a stabilizing gate, preventing oleic acid from exiting the active site and ensuring its stability within the core. This selective stabilization of oleic acid, while allowing the partial unbinding of diolein, also suggests a dual function for Cor a 8 as a moonlight enzyme, capable of catalyzing the hydrolysis reaction and transporting single-chain lipids while releasing the complementary part.

It is interesting that the same amino acids possibly involved in the hydrolysis are responsible for the oleic acid stability. This mechanism could be necessary to restrain the enzyme activity until the oleic acid is delivered, similarly to the aconitase moonlighting enzyme that needs to lose the Fe-S cluster to become the iron responsive element (IRP1) [52], [53].

Finally, our phylogenetic analysis conducted on all land plants highlighted that the catalytic amino acids (Tyr103, Arg63) appeared and are conserved only within the angiosperms. This result strongly suggests that the protein may have acquired this new enzymatic activity to play an essential role in both anabolism and catabolism of lipids in oilseed plants. Deciphering its unique catalytic properties, as well as understanding its interactome, will be a crucial step in developing strategies to regulate lipid-related processes, with significant implications for the food industry as well as possible novel biotechnological applications.

## Author contributions

S.A. conceived and supervised the project. A.F. and G.D.N. purified the protein and carried out the kinetics and FESEM analysis. E.S. performed bioinformatic analyses. M.L.D.S. and F.F. carried out the molecular dynamics. V.S. and F.D.P. performed the mass spectrometry analysis. M.M and S.O.B. performed the TLC experiment. G.V. and A.C. helped with establishing the protein purification protocol. F.P. carried out the elaboration of enzyme kinetics parameters. G.G. supplied fresh hazelnuts and helped with the experimental design. M.Mz. guided in carrying out the electro-microscopy measurements. A.G.B and S.I. carried out the secondary structure characterization.

## Acknowledgments

We would like to acknowledge Prof. Chiara Cordero and Prof. Franco Bonomi for the much-appreciated scientific discussion. This work was supported by Soremartec S.R.L. (Industrial_Grants ADIS_CT_RIC_22_01, ADIS_CT_RIC_22_02) and by University of Turin (Ricerca Locale Grants ADIS_RILO_21_01, OLIS_RILO_21_02, MARM_RILO_22_05.

