## Supplementary data for "*Corylus avellana* non-specific lipid-transfer protein Cor a 8 is a moonlighting enzyme with a new lipase activity"

FIGURE SUPPLEMENTARY 1

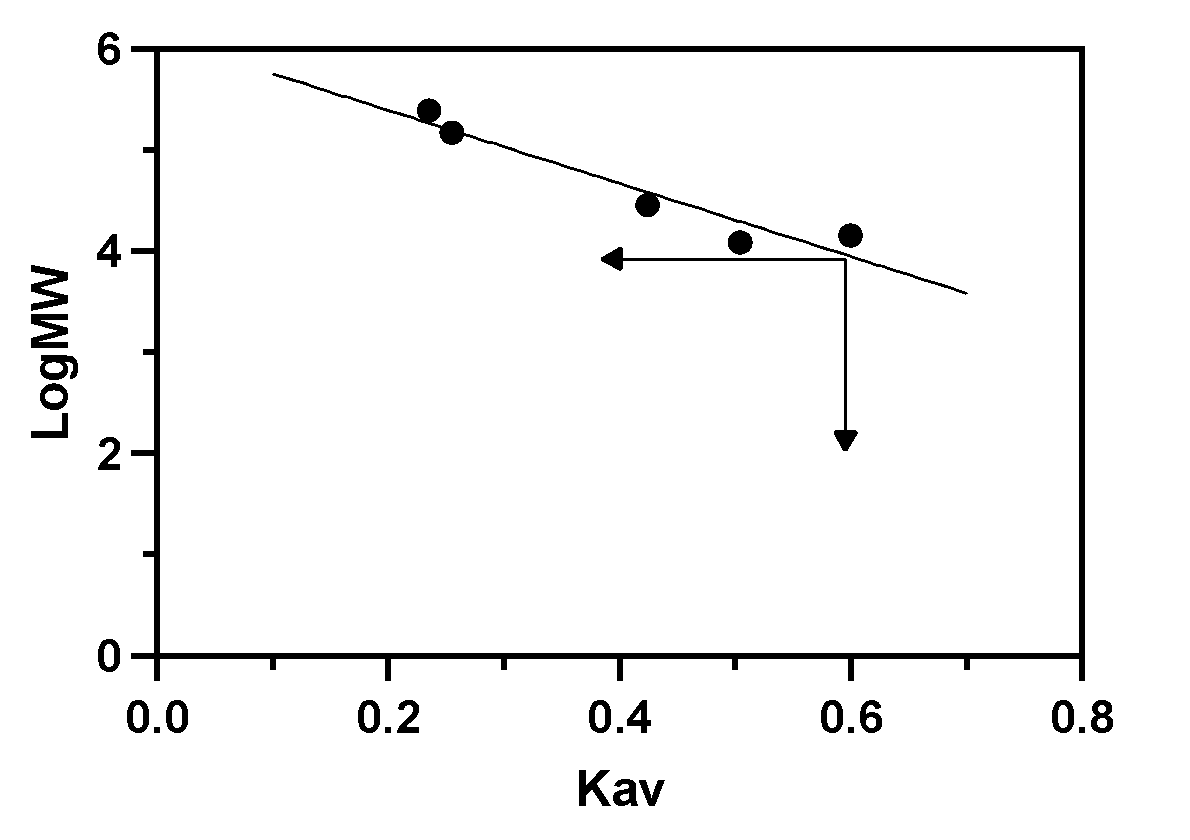

Figure S1 – **Calibration curve for apparent molecular mass of Cor a 8 in native condition.** SEC mobility Kav is reported in function of LogMW, from smaller to the higher Kav were used the following calibration standards: Catalase from bovine liver, Alcohol Dehydrogenase from Saccharomyces cerevisiae, Carbonic Anhydrase from bovine erythrocytes, Cytochrome C from horse heart muscle, Lysozyme from chicken egg white. The double black arrow identifies the Kav (0.595) and corresponding LogMW (3.966) value of the purified lipase via anionic exchange chromatography.

FIGURE SUPPLEMENTARY 2

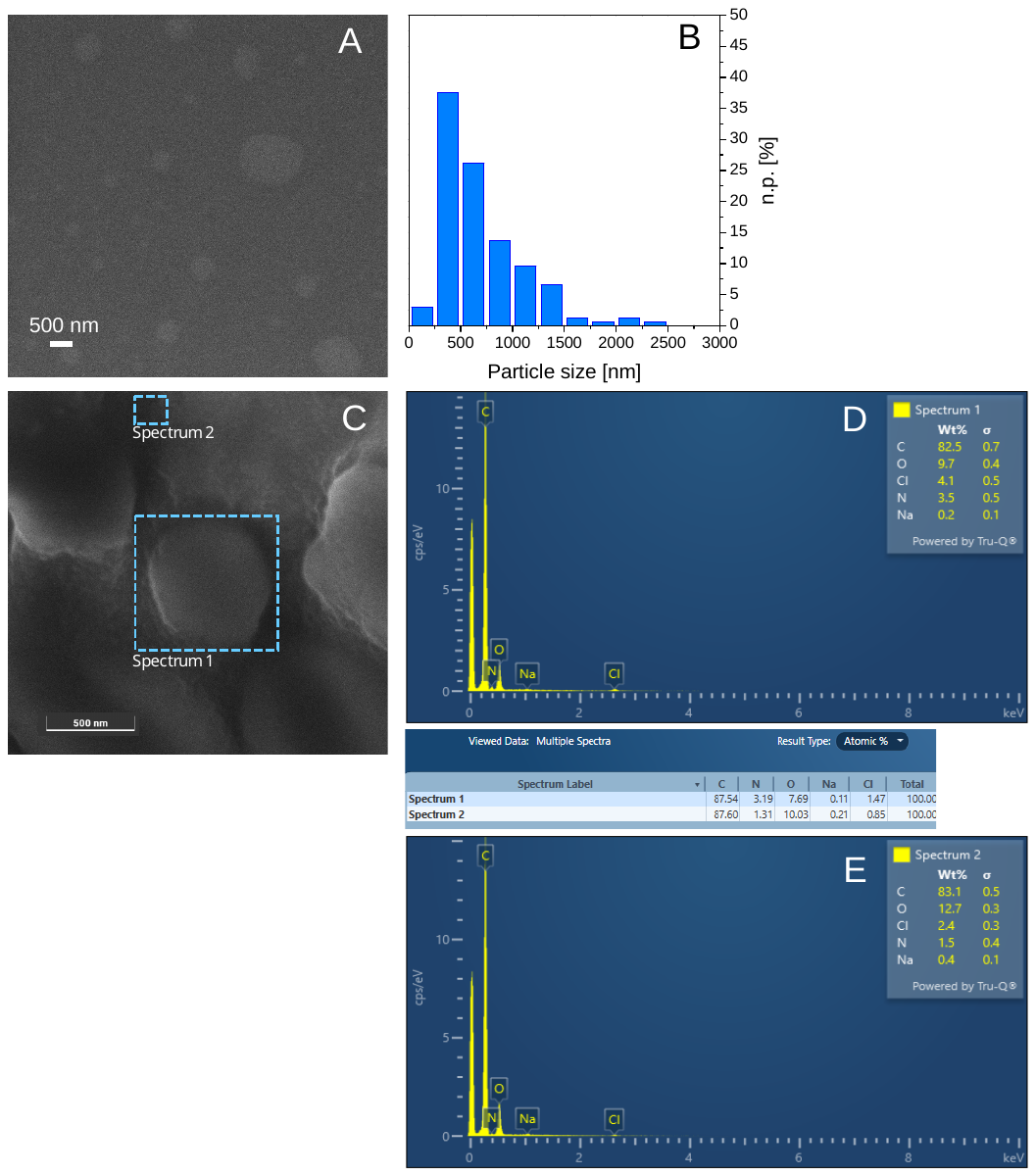

Figure S2 – **Field Emission Scanning Electron Microscopy (FESEM) of the lipid nanoparticles stabilized by Triton X-100 and in the presence of the pNPL substrate** – FESEM representative images (A and C) and particle size distribution (B). EDS spectra of the regions highlighted by the dashed boxes in C (D and E). Images collected at 5 keV with the in-beam cross-free mode SE detector. n.p.[%] represents the number of counted particles of diameter d_i_. Instrumental magnification: 15000× and 130000×, respectively. Dashed blue boxes highlight the region of sample where the EDS spectra where acquired, spectrum 1 a micelle, spectrum 2 micelle free region as a reference control.

FIGURE SUPPLEMENTARY 3

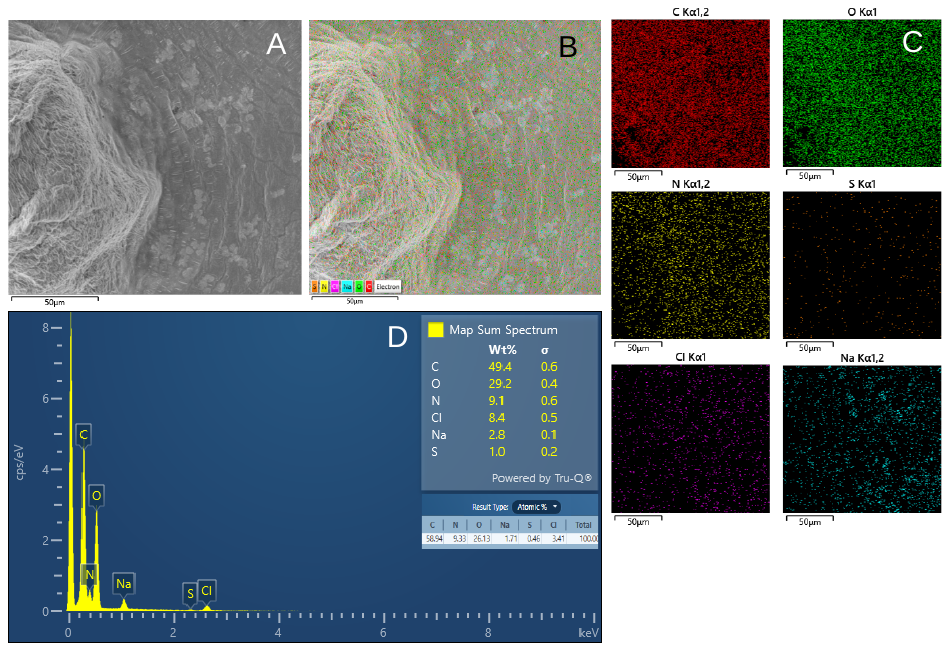

Figure S3 – **Field Emission Scanning Electron Microscopy (FESEM) of the enzyme and lipid nanoparticles in the presence of the pNPL substrate** - FESEM representative image (A) and EDS layered image (B). EDS maps for C, O, N, S, Cl and Na (C) and EDS sum spectrum of the region shown in A (D). Image collected at 5 keV with the in-beam SE detector. Instrumental magnification: 1200×.

FIGURE SUPPLEMENTARY 4

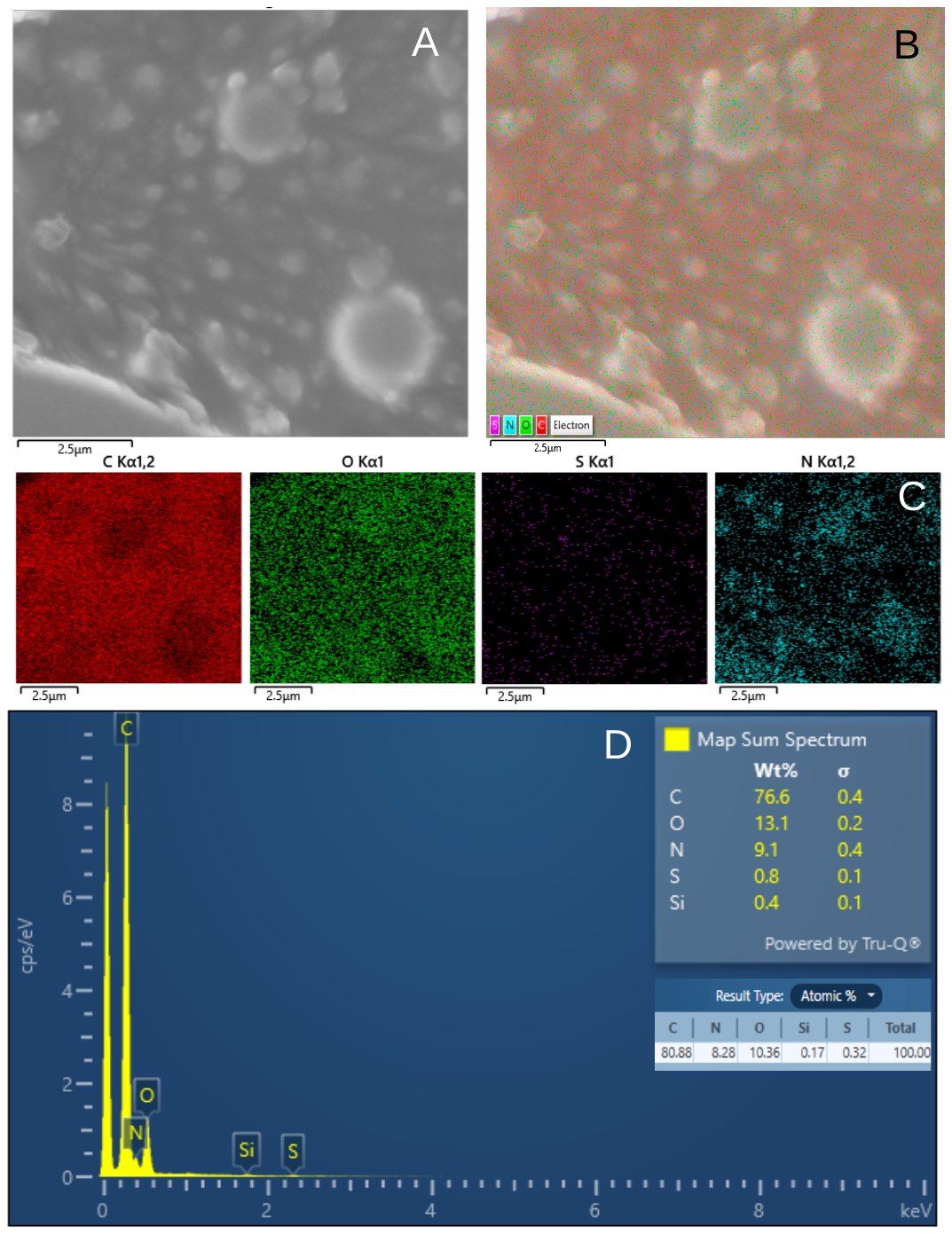

Figure S4– **Field Emission Scanning Electron Microscopy (FESEM) of the enzyme in the presence of Triton X-100** - FESEM representative image (A) and EDS layered image (B). EDS maps for C, O, S and N (C) and EDS sum spectrum of the region shown in A (D). Image collected at 5 keV with the in-beam SE detector. Instrumental magnification: 21000×.

FIGURE SUPPLEMENTARY 5

The reversible Michaelis-Menten (RMM) scheme is the simplest set of reactions to describe the catalyzed transformation of a substrate S into a product P that also consider the formation of an intermediate complex. The RMM scheme reads as follows:

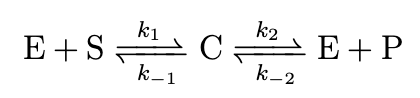

(1)

Let us indicate with C_α_ (α = E,S,C,P) the concentrations of different reactants. The rate equations associate with the chemical scheme (1) read

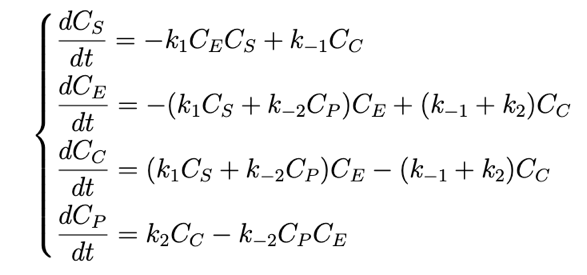

(2)

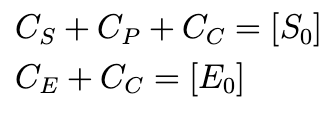
Of course, only two of the rate equations (2) are independent, in view of the two mass conservation laws

(3)

where [*S*_0_] and [*E*_0_] are the total substrate and enzyme concentration, respectively. We choose *C_S_* and *C_P_* as independent variables. Let us introduce the following non-dimensional concentrations and parameters

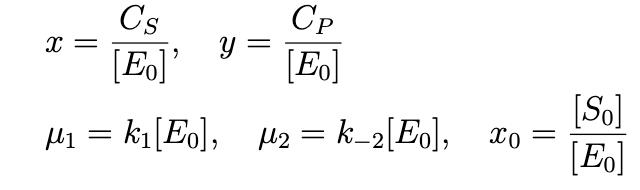
(4)

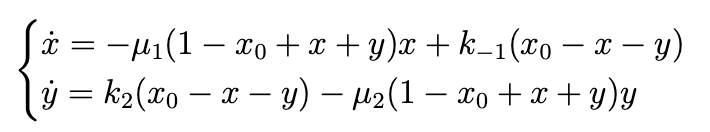
Recalling also the two conservation laws (3), the two independent rate equations take the following form

(5)

In our fitting script, we use the differential evolution algorithm [54] to minimize a cost function. A call to the cost function routine computes the least-squares deviation between the data (all the available measurements of *C_P_*(*t_i_*) for all time points *t_i_* and substrate concentrations [*S*_0_]) and the theory. The latter is computed by performing a numerical integration of Eqs (5) with the LSODE (Livermore Solver for Ordinary Differential Equations) algorithm as implemented in the Scipy function Odeint, corresponding to a given choice of the free parameters in the model and with initial conditions *x*(0) = *x*_0_, *y*(0) = 0.

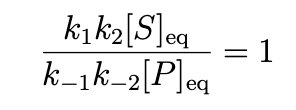
 Since we do not know how much of the purified enzyme is active in the solution, we treat [E_0_] as a floating parameter, with the meaning of the concentration of active enzyme. Overall, therefore, there are five free parameters in our RMM model, the four rates *k*_1_, *k*_−1_, *k*_2_, *k*_−2_ and the concentration [*E*_0_]. We found that far better fits could be obtained if the rate *k*_−2_ was computed from the detailed balance constraint as a function of the ratio between the equilibrium product and substrate concentrations. At thermodynamic equilibrium, detailed balance should hold, namely

(6)

Hence, our choice of the five floating parameters was

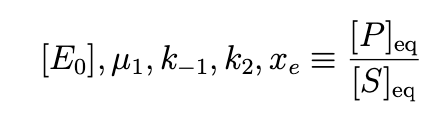
(7)

while *k*_−2_ was computed from the parameters (7) via Eq. (6).

Differential evolution is an evolutionary fitting algorithm suitable for solving complex global optimization problems by progressively refining a potential solution through an iterative evolutionary process. Since it is intrinsically stochastic in nature and does not require initial guesses for the free parameters (only bounds), the results of many independent optimization runs can be used to assess the statistical variability of the best-fit results. In Fig. S5, we illustrate the results of this statistical analysis. It is apparent that all the best-fit values are distributed around their means (those tabled in the main text) with a spread, measured in units of standard deviation over mean, that is at most 4 %.

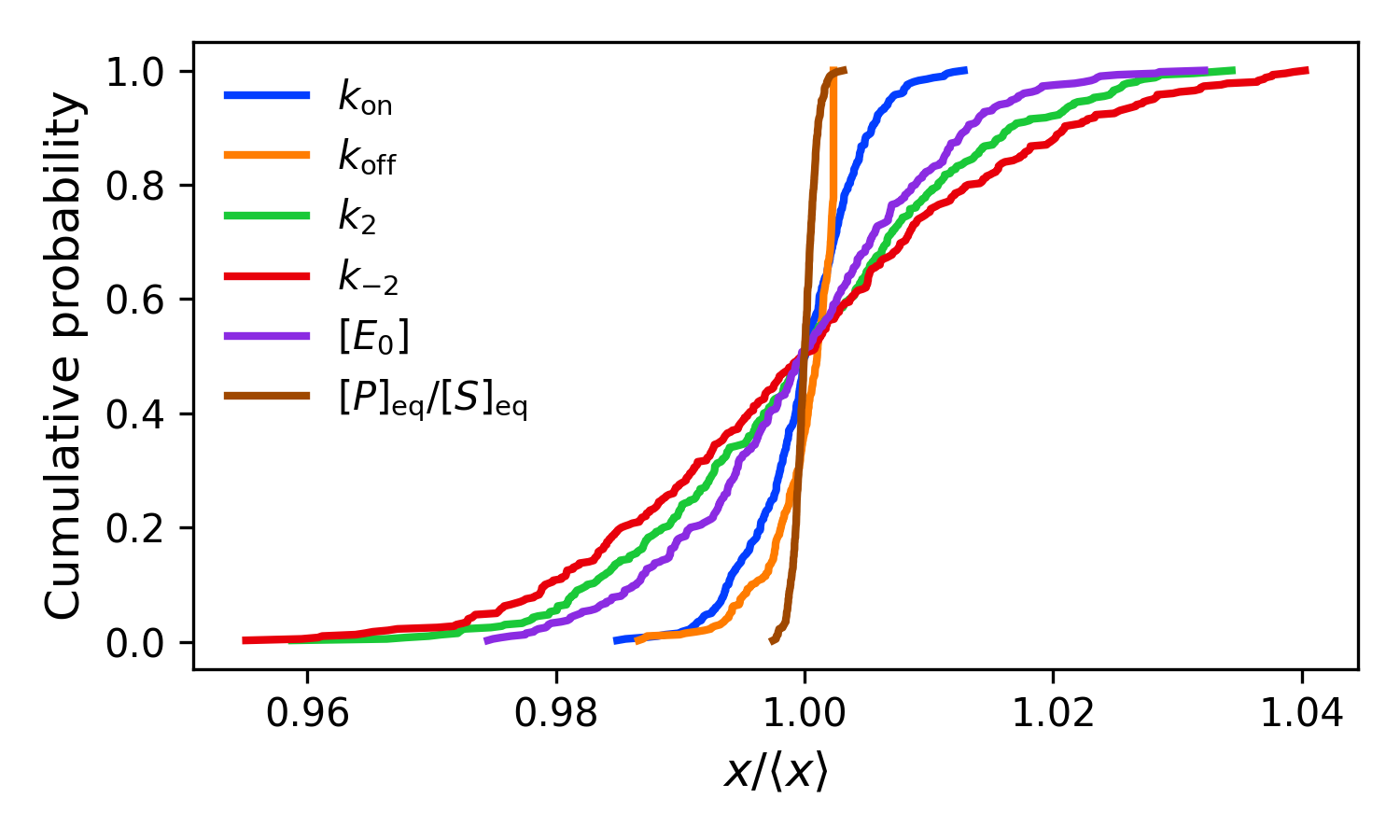

Figure S5 **Accuracy of the fitting procedure.** Cumulative probabilities of the best-fit values of the five free parameters (and the corresponding values of *k*_−2_) obtained for *N_R_* = 400 independent minimization runs of the enzyme kinetics. The cumulatives (i.e. the probability that a parameter is greater than a fixed value, also called distribution functions) have been computed by normalizing the parameters (*x* = [*k*_on_, *k*_off_, *k*_2_, *k*_-2_, [*E*_0_], [*P*]_eq_/[*S*]_eq_]) through their average values computed over the 400 minimization runs..^[1]^

Figure S6

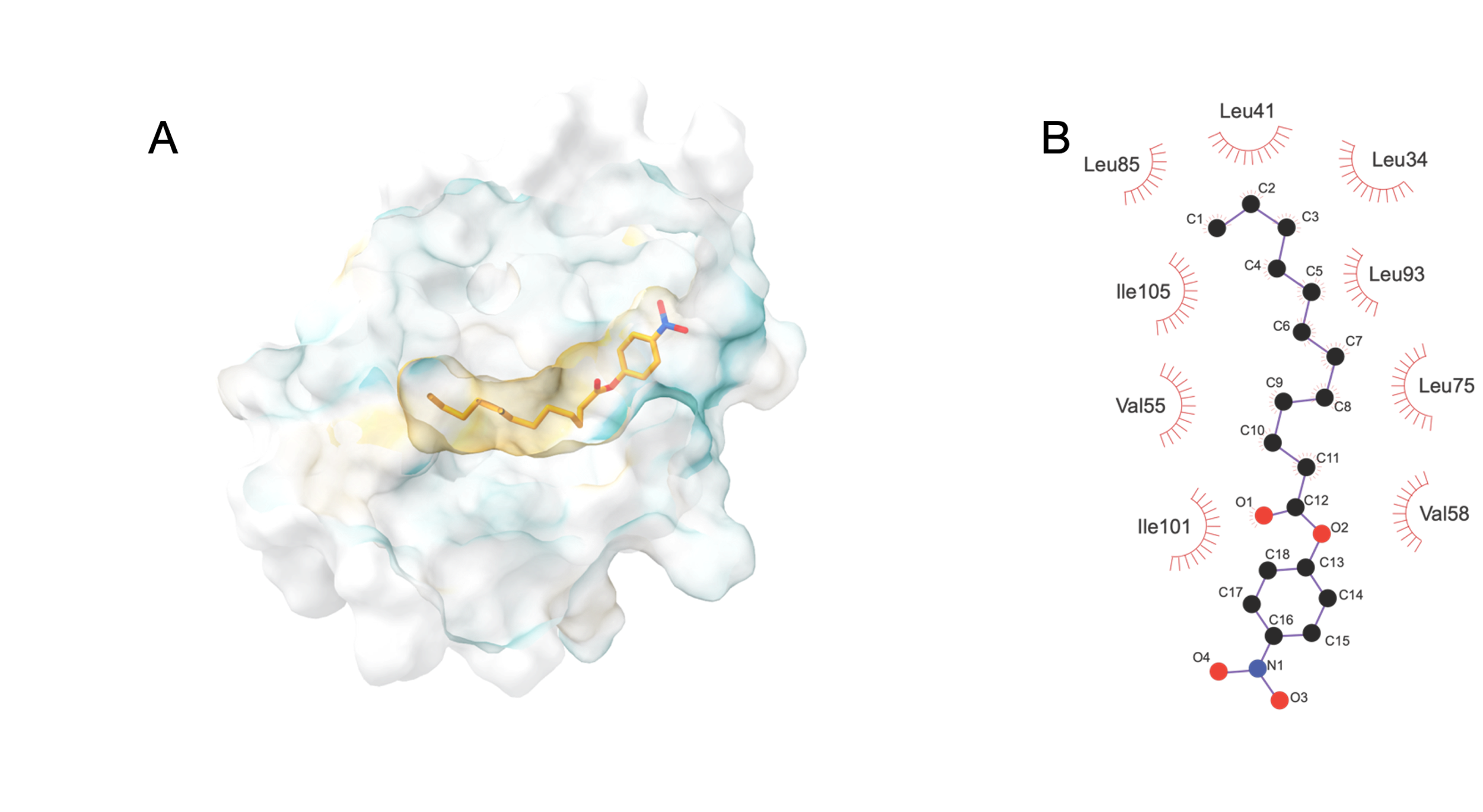

Figure S6. **pNLP docking in Cor a 8**. A) The pNPL modelled in the hydrophobic cavity. Surface is shown in the Kyte-Doolittle scale, with blue representing the most hydrophilic white at neutral and orange-red for the most hydrophobic. The lipid is shown as sticks. Figure was created using UCSF Chimera version 1.7. B) A representation of the hydrophobic interaction is shown by LigPlot+ [55].

FIGURE SUPPLEMENTARY 7

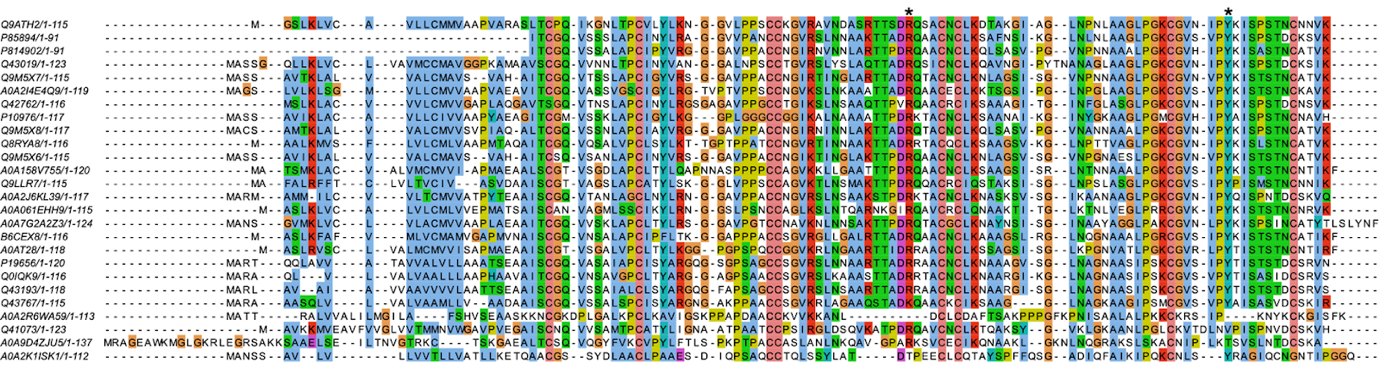

Figure S7 - **nsLTP sequence alignment with Clustal and visualized in Jalview**. The proteins are labelled by the respective UniProtID code (Table 1). The Cora8 corresponding Arg68 and Tyr103 are highlighted with an asterisk.

TABLE S1

| **Species** | **UniProt ID** |
| --- | --- |
| *Adiantus capillus-veneris* | A0A9D4ZJU5 |
| *Arabidopsis thaliana* | Q9LLR7 |
| *Arachis hypogaea* | B6CEX8 |
| *Corylus avellana* | Q9ATH2 |
| *Gossypium hirsutum* | Q42762 |
| *Hevea brasiliensis* | Q8RYA8 |
| *Hordeum vulgare* | Q43767 |
| *Juglans regia* | A0A2I4E4Q9 |
| *Lactuca sativa* | A0A2J6KL39 |
| *Lens culinaris* | A0AT28 |
| *Macadamia integrifolia* | A0A7G2A2Z3 |
| *Malus domestica* | Q9M5X7 |
| *Marchantia polymorpha* | A0A2R6WA59 |
| *Morus nigra* | P85894 |
| *Oryza sativa* | Q0IQK9 |
| *Physcomitrium patens* | A0A2K1ISK1 |
| *Pinus taeda* | Q41073 |
| *Pisum sativum* | A0A158V755 |
| *Prunus avium* | Q9M5X8 |
| *Prunus dulcis* | Q43019 |
| *Prunus persica* | P814902 |
| *Pyrus communis* | Q9M5X6 |
| *Sorghum bicolor* | Q43193 |
| *Spinacia oleracia* | P10976 |
| *Theobroma cacao* | Q9M5X7 |
| *Zea mays* | P19656 |

Table S1. Species and UniProt IDs in Figures 7A and 7B
